# Hidden neural states underlie canary song syntax

**DOI:** 10.1101/561761

**Authors:** Yarden Cohen, Jun Shen, Dawit Semu, Daniel P. Leman, William A. Liberti, L. Nathan Perkins, Derek C. Liberti, Darrell Kotton, Timothy J. Gardner

## Abstract

Coordinated skills such as speech or dance involve sequences of actions that follow syntactic rules in which transitions between elements depend on the identity and order of past actions. Canary songs are comprised of repeated syllables, called phrases, and the ordering of these phrases follows long-range rules, where the choice of what to sing depends on song structure many seconds prior. The neural substrates that support these long-range correlations are unknown. Using miniature head-mounted microscopes and cell-type-specific genetic tools, we observed neural activity in the premotor nucleus HVC as canaries explore various phrase sequences in their repertoire. We find neurons that encode past transitions, extending over 4 phrases and spanning up to 4 seconds and 40 syllables. These neurons preferentially encode past actions rather than future actions, can reflect more than a single song history, and occur mostly during the rare phrases that involve history-dependent transitions in song. These findings demonstrate that network dynamics in HVC reflect preceding behavior context relevant to flexible transitions.

## Main body

Flexible behavior often contains complex transitions - points where the next action depends on memory of choices made several steps in the past. Canary song provides a highly tractable model for studying flexible sequence production. These songs are highly complex, but are composed of well-defined units or syllables. These syllables are produced in trilled repetitions known as phrases (Figure 1a) that are approximately one second long. Phrases are organized in sequences to produce songs that are typically 20-40 seconds long, and the sequences of phrases exhibit long-range syntax rules^1^. Specifically, phrase transitions following about 15% of the phrase types show strong context dependence that extends 2-5 phrases into the past. These long-range correlations extend over dozens of syllables, spanning time intervals of several seconds (Figure 1b,c). In premotor brain regions, neural activity supporting long-range complex transitions will reflect context information as redundant representations of ongoing behavior^2–5^. Such representations, referred to here as ‘hidden neural states’, are predicted in models of memory-guided behavior control^6^, but are challenging to observe during unconstrained motion in mammals^7–14^ or in songbirds with simple syntax rules^15^.

**Figure 1.**
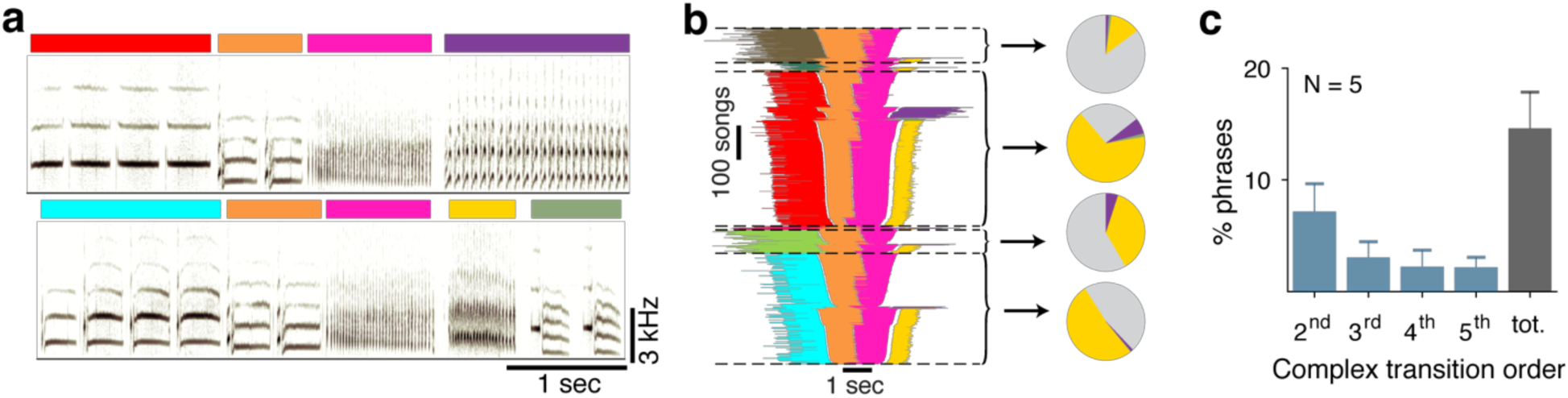
Long range syntax rules in canary song. **a.** Two example spectrograms of Canary song. The colored bars indicate the identity of the phrase, which is composed of basic elements called syllables. Both of these examples contain a common phrase transition (orange to pink) but differ in the preceding and following phrases. **b.** A summary of all phrase sequences that contain this common transition reveals that the choice of what to sing after the pink phrase depends on the phrases that were produced earlier. Lines represent phrase identity and duration. Song sequences are stacked (vertical axis) sorted by the identity of the 1^st^ phrase, the last phrase and then the center phrases’ duration. Pie charts show the frequency of phrases that follow the pink phrase, calculated in the subset of songs that share a preceding sequence context (separated by dashed lines). Grey represents the song end, and other colors represent a phrase pictured in the first panel. The pink phrase precedes a ‘complex transition’, the likelihood that a particular phrase will follow it is dependent on transitions several phrases in the past. **c.** Percent of phrases that precede complex transitions of different orders. Bars and error bars show mean and SE in 5 birds.

Like motor control in many vertebrate species, canary song is governed by a ‘cortico-thalamic loop’ that includes the premotor nucleus HVC^16–18^. In stereotyped songs of zebra finches, HVC projection neurons produce stereotyped bursts of activity time-locked to song^17^. This neural precision drives reliable motor outputs and relays timing reference to the basal ganglia^19^, integral to both cortical^7,20^ and striatal^21^ mechanisms of sequence generation^22–27^. In the more variable syllable sequences of Bengalese finches, the same projection neurons fire differently depending on the neighboring syllables^15^, supporting sequence generation models that include hidden states^6^. However, the time-frame of the song-sequence neural correlations in that seminal experiment was relatively short (under 100ms). In contrast, long-range correlations in human syntax can extend for tens of seconds and beyond and exhibit long-order rules. At present it is not known if redundant premotor representations in songbirds can support working memory for syntax control over timescales longer than 100ms and whether this activity is phasic^28^, or takes sustained^8^ or ramping forms. To further dissect the mechanisms of working memory for song, we examined motor control of song in the canary, *serinus canaria*, a species with long-range syntax rules^1^.

We used custom head mounted miniature microscopes to record HVC activity during the song production in freely moving canaries (Figure 2b). In repeating sequences, spanning up to four phrases, we found that individual neurons are active differently depending on the identity of previous, or upcoming non-adjacent phrases - reflecting phrase-locked hidden network states encoding song ‘context’ beyond the ongoing behavior. The context-dependent changes in canaries extend over 4000ms, demonstrating a deep many-to-one mapping between HVC state and song syllables in this species. We find that neural activity correlates more often with the song’s past than its future, and that complex, context-dependent neural activity occurs selectively in complex parts of the behavior, where phrases are followed by context-dependent transitions. Additionally, we find that HVC neurons can be selective to a single song context or exhibit mixed selectivity – showing strong activity dependent on multiple song histories. Together, these findings reveal a previously un-described pattern of neural dynamics that can support structured, context-dependent song transitions and validate predictions of long-range syntax generation by hidden neural states^6,29^ in a complex vocal learner.

**Figure 2.**
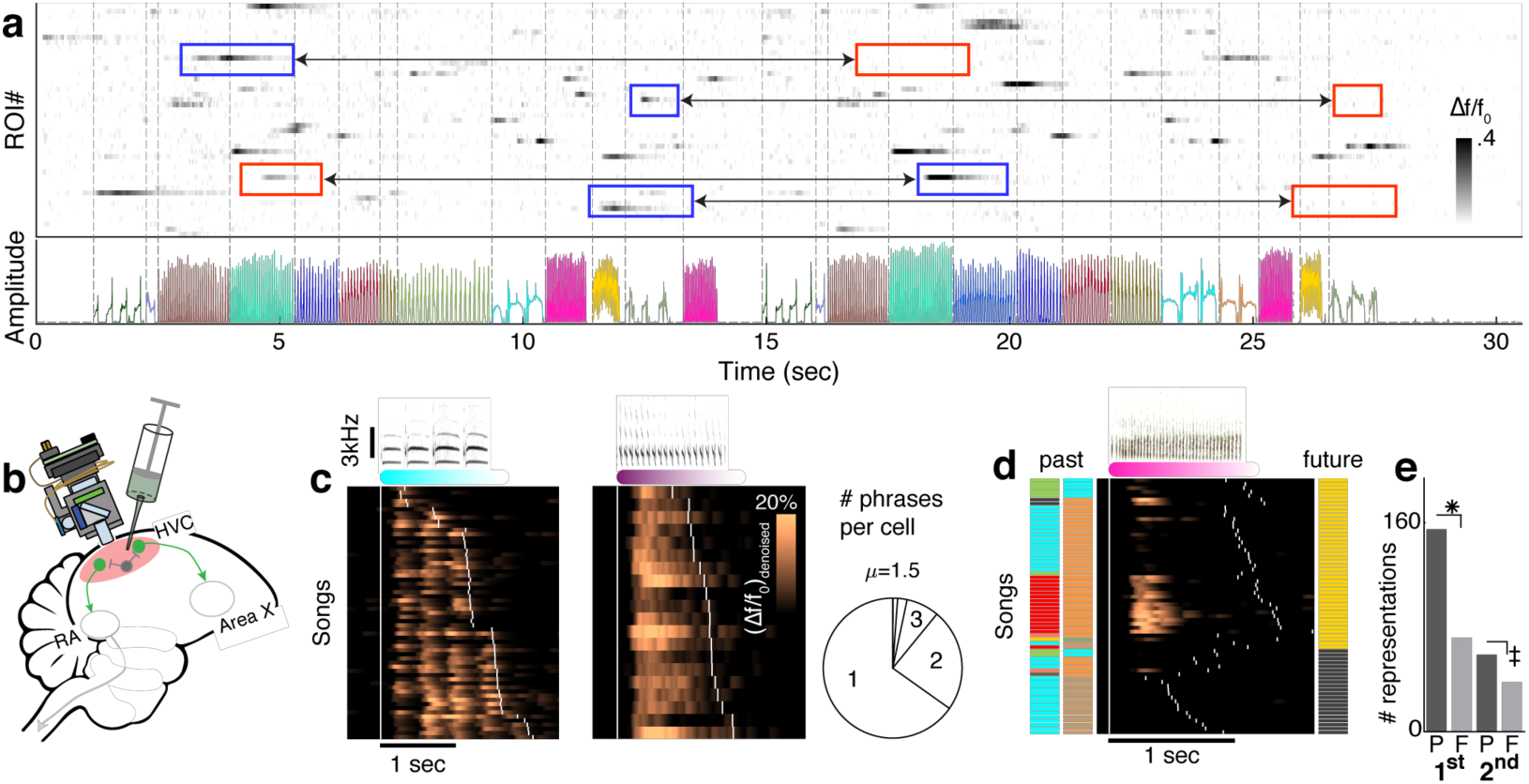
HVC projection neuron activity reflects long-range information about phrase transitions. **a.** Fluorescence (*Δf*/*f*_0_) of multiple ROIs during a singing bout reveals sparse, phrase-type-specific activity. Phrase types are color coded in the audio amplitude trace, and dashed lines mark phrase onsets. Context-dependent cells show larger phrase-specific signal in one context (blue frames) than another (connected red frames). **b.** Experimental paradigm. Miniature microscopes were used to image GCaMP6f-expressing neurons in HVC, transduced via lentivirus injection. **c.** Most cells are phrase-type-specific. Neural activity is aligned to the onset of phrases with long (left) and short (right) syllables. Songs (y-axis) are sorted by the phrase duration, and white ticks indicate phrase onsets. Pie shows fractions of neurons that are active during just one, two or three separate phrase types (methods). **d.** ROI activity during a phrase that is strongly related to 2^nd^ upstream phrase identity. Neural activity is aligned to the onset of the current phrase. Songs are arranged by the ending phrase identity (right, color patches), then by the phrase sequence context (left, color patches), and then by duration of the pink phrase. White ticks indicate phrase onsets. **e.** More neural representations reflect past events than future events. 258 different ROIs had a total of 324 significant correlations with adjacent (1^st^ order, 2 left bars) and non-adjacent (≥2^nd^ order, 2 right bars) phrases. The correlations are separated by phrases that precede (P) or follow (F) the phrase, during which the signal is integrated. Binomial z-test evaluate significant differences (*: p < 1e-10, ‡: p < 0.002).

## Results

### Canary songs are characterized by structure on three time-scales: syllables, phrases, and phrase sequence syntax

Inspired by the success of machine learning methods for human speech recognition^30^, we developed a song segmentation and annotation algorithm that automated working with large datasets (>5000 songs from 3 birds; Supplementary Figure 1 - 1, methods). The birds’ repertoire included 24-37 different syllables with typical durations between 10-350 msec (Supplementary Figure 1 - 2c,d). The average number of syllable repeats per phrase type ranged from 1 to 38 with extreme cases of individual phrases exceeding 10 seconds and 120 syllables (Supplementary Figure 1 - 2a,b). Transitions between phrases can be completely deterministic, where one phrase type always follows another, or flexible, where multiple phrase types can follow a given phrase (sonograms in Figure 1a, stack in Figure 1b). Phrase transitions are well identified by the change of the repeated syllable with no confusion by other sequence components - within-phrase syllable acoustics, phrase and inter-phrase gap durations (Supplementary Figure 1 - 3 and Supplementary Figure 1 - 5 illustrate this for single phrase sequences and syllable repertoires). In very rare cases, transitions can contain an aberrant syllable that cannot be stably classified (c.f. Supplementary Figure 1 - 6).

In the following analysis, we focused on neural activity that is correlated with phrase sequence variability. In our dataset, 95% of all phrases are trills of multiple syllables and only 6.1% of those are shorter than the decay time constant of the calcium indicator we used (GCaMP6f, 400 msec^31^, Supplementary Figure 1 - 2f). As in finches, we found that HVC projection neuron activity in canaries was sparse in time. This, combined with the long phrase duration (Supplementary Figure 1 - 2d), allowed us to resolve phrase level sequence correlations with little or no limitations resulting from the calcium indicator’s temporal resolution.

### A small set of phrase types precede complex transitions

To investigate long-range syntax rules in canary song we examined the context dependence of phrase transitions. As shown previously in another strain of canaries^1^, we find that a small subset of phrase types precede ‘complex’ transitions - behavioral transitions that depend on the multi-step context of preceding phrases. Specifically, the probability of transition outcomes can change by almost an order of magnitude depending on the identity of the 3 preceding phrases (Figure 1b). Such song context dependence is captured by a 3^rd^ order Markov chain. Supplementary Figure 1 - 4 shows the significant long-range context-dependent transitions for two birds.

### HVC neural activity reflects long-range sequence information

To characterize the neural activity supporting complex transitions, we imaged neurons that expressed the genetically-encoded calcium indicator GCaMP6f in freely-behaving adult male canaries (Serinus canaria, n=3, age > 1yr, recording in left hemisphere HVC^16^). The indicator is selectively expressed in excitatory neurons (Supplementary Figure 2 - 1). With this technique, we are able to record neural activity via fluorescence dynamics extracted from annotated regions of interest (ROIs, Supplementary Figure 2 - 4, see methods).

Fluorescence signals in these ROIs was sparse in time, and time-locked to specific syllables or phrases, which is consistent with findings in other species with simpler songs^15,17^ (Figure 2a,c, Supplementary Figure 2 - 6). Out of N = 2010 daily annotated ROIs from 3 birds (35 +/-15, mean +/-SD ROIs per animal per day), about 90% are active in just one or two phrase types. Additionally, we find that the pattern of phrase-locked activity in some ROIs changes depending on song context. For example, some ROIs showed weaker or no activity in one song context while demonstrating strong activity in another song context (Figure 2a).

Importantly, we find that this context-dependent activity is strongly influenced by the identity of non-adjacent phrases. For example, Figure 2d shows the de-noised fluorescence signal raster from a ROI, locked to the phrase type marked in pink, displaying a dramatic variation in activity ((*Δf*/*f*_0_)_*denoised*_, methods) depending on the 2^nd^ phrase in the sequence’s past – a 2^nd^ order correlation. This sequence preference is quantified by integrating the ROI-averaged signal (Supplementary Figure 2 - 2a, 1-way ANOVA, p < 1e-8, Supplementary Figure 2 - 3g,h show the source ROI). We find ROIs with signals that relate to the identity of past and future non-adjacent phrase in all 3 birds (Supplementary Figure 2 - 3). Across all animals, 18.2% of the daily annotated ROIs showed sequence correlations extending beyond the current active syllable. 15% had 1^st^ order correlations where activity during one phrase depends on the identity of an adjacent phrase, and 5.6% had ≥2^nd^ order relations (Figure 2d, Supplementary Figure 2 - 9).

These sequence dependencies could potentially be explained by other factors inherent to the song that may be more predictive of phrase sequence than HVC activity. For example, transition probabilities following a given phrase could potentially depend on the phrase duration^1^, on the onset and offset timing of previous phrases, and on the global time since the start of the song – implicating processes such as neuromodulator tone, temperature buildup, or slow adaptation to auditory feedback^32–37^ (c.f. Supplementary Figure 2 - 7a,b, Supplementary Figure 2 - 8). To rule out these explanations, we used multivariate linear regression and repeated the tests for sequence-correlated neural activity after discounting the effect of these variables on the neural signals. We found that 31.9% (31/97 from 3 birds) of ≥2^nd^ order relations and 56.8% (129/227 from 3 birds) of the 1^st^ order relations remain significant (Supplementary Figure 2 - 7c, Supplementary Figure 2 - 2).

The sequence-correlated ROIs tend to reflect past events more often than future events. Out of N = 324 significant phrase sequence-neural activity correlations, 66% reflect preceding phrase identities (p < 1e-10, binomial z-test). This bias is also found separately in 1^st^ or higher order correlations (Figure 2e, 68.2% and 60.8% respectively. Both percentages are significantly larger than chance, p < 1e-10, p < 0.002, binomial z-test). This bias persists if we consider ROIs that overlap in footprint and sequence correlation across days as a unified representation (Appendix A).

These findings suggest that for a subset of HVC neurons, calcium signals are not just related to present motor actions, but convey the context of past events across multiple syllables.

### Sequence-correlated ROIs reflect preceding phrase identities up to four steps apart

To produce songs with long-range syntax rules, a memory of previous elements sung must influence the current syllable choice. It is not known how this information about past choices is carried forward in time during canary song. Clues for such a process can be seen in one example where, in a fixed sequence of four phrases, we found ROIs that carry information about the identity of an early phrase during each phrase in the following sequence (Figure 3a,b, Supplementary Figure 3 - 3a, 1-way ANOVA showing significant modulation of neural activity with the identity of the past phrase). In this example, the ROIs that reflect long-range information continue to do so even if the final phrase in the sequence is replaced by the end of song, suggesting that their activity reflects prior song context rather than some upcoming future syllable choice. (Supplementary Figure 3 - 3b, 1-way ANOVA, p < 5e-6 and p < 0.08 for ROIs 50 and 36, when replacing the last phrase with the end of song). This example suggests that a chain of neurons reflecting “hidden states” or information about past choices could provide the necessary working memory for complex phrase transition rules.

**Figure 3.**
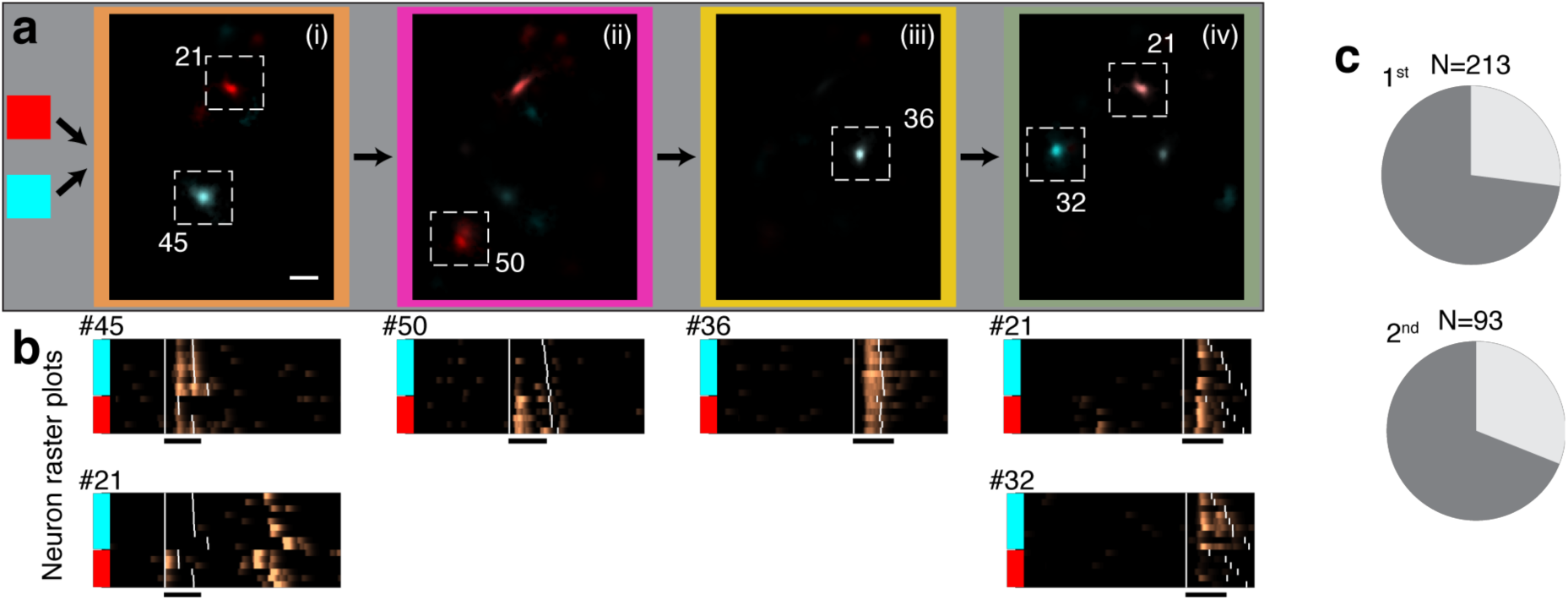
Sequence-correlated HVC neurons reflect preceding context up to four phrases apart and aggregate around context-dependent transitions. **a.** A sequence of four phrases (i-iv, color coded) is preceded by two upstream phrase types (red or cyan). Average maximum projection denoised images (methods) are calculated in each sequence context during each phrase in the sequence (i-iv) and overlaid in complementary colors (red, cyan) to reveal context-preferring neurons. **b. (***Δf*/*f*_0_)_*denoised*_ rasters for the ROIs in panel (a). Songs are ordered by the preceding phrase type (colored bars). Supplementary Figure 3 - 1a shows the statistical significance of song context relation. **c.** Fraction of sequence-correlated neurons found in complex transitions (e.g. in Figure 1). Pie charts separate 1^st^ order and higher order (≥2^nd^) sequence correlations. Dark grey summarizes the total fraction for two birds.

### Sequence-correlated ROIs are predominantly found during context-dependent transitions

The phrases in Figure 3 are phrase types that lead to complex transitions or directly follow them (in Figure 1). If HVC neurons are involved in driving phrase transitions that follow long-range syntax rules, then they should represent song context information predominately around complex behavior transitions, when such information is needed to bias transition probabilities. Accordingly, at the population level, we find more sequence-correlated ROIs around complex transitions; about 70% of sequence-correlated ROIs are found during the rare phrase types that participate in complex transitions (Figure 3c, Supplementary Figure 3 - 3). This bias persists if we consider ROIs that overlap in footprint and sequence correlation across days as a unified representation (Appendix A). Separating representation of past context and future action we find that, in complex transitions, ROIs predominately represent the identity of a preceding phrase (Supplementary Figure 3 - 3a,b, multi-way ANOVA test effects of preceding and following phrases, revealing a 3:1 bias in representing past context in complex transitions. p < 1e-13, binomial z-test). This bias does not occur outside of complex transitions (Supplementary Figure 3 - 3c, p > 0.2 binomial z-test), suggesting that neural coding for past context is dominant in transitions depending on this information.

### Sequence-correlated ROIs can prefer more than one song context

The concentration of sequence-correlated ROIs in complex transitions suggests that HVC neurons can encode behaviorally relevant song contexts. Examining the song contexts preferred by single ROIs, we find that some are selective to a single prior context, i.e., their mean activity, following one context is significantly larger than for all others (methods). Additionally, many ROIs have a clear preference to more than a single past (Figure 4a,b, Supplementary Figure 4 - 1). Among all context-selective ROIs, 19% and 14% have such mixed context selectivity in 1^st^ order and ≥ 2^nd^ order sequence correlations respectively. Additionally, 44% and 48% of the context-selective ROIs strictly prefer one past out of several contexts in 1^st^ order and ≥2^nd^ order correlations (Figure 4c, from 3 birds).

**Figure 4.**
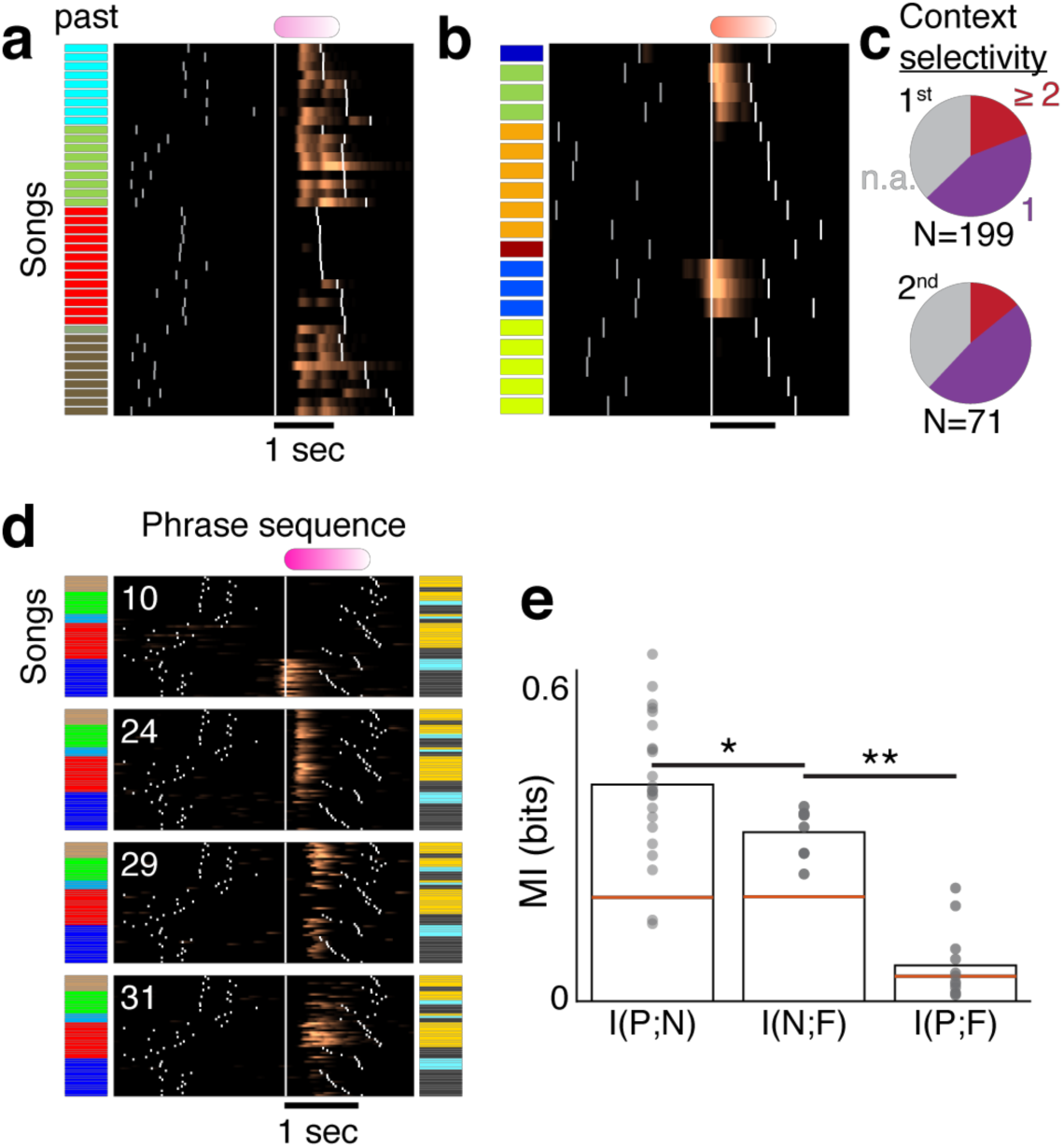
HVC neurons can be tuned to distinct, and complementary, preceding contexts. **a**,**b.** ROIs can respond to multiple preceding contexts. (*Δf*/*f*_0_)_*denoised*_ traces are aligned to a specific phrase onset, arranged by identity of preceding phrase (color barcode). White ticks indicate phrase onsets. **c.** Context-correlated ROIs significantly prefer one (purple) or more (red) song contexts. Pies separate ROIs with selectivity to the 1^st^ (top) and 2^nd^ (bottom) preceding phrases. **d.** Four example jointly-recorded ROIs that exhibit complementary context selectivity. Color bars indicate the phrase identities preceding and following a fixed phrase (pink). For each ROI (rasters), (*Δf*/*f*_0_)_*denoised*_ traces are aligned to the onset of the pink phrase (x-axis, bar marks 1 sec) arranged by the identity of the preceding phrase, by the following phrase and finally by the duration of pink phrase. White ticks indicate phrase onsets. **e.** The network holds transition-relevant context information in panel (d). Mutual information between the identity of past (P) and future (F) phrase types (right bar. Variables match barcode in panel d) is significantly smaller than the information held by the network states about the past and future contexts (left bars. N is the 4-ROIs activity). Dots mark bootstrap assessment shuffles. Bars mark the mean. Red lines mark the 95% level of the mean in shuffled data (methods). *: p<1e-4, **: p< 1e-6, bootstrapped z-test.

### Jointly-recorded ROIs predict behavior prior to a complex transition

Neurons with distinct song context preferences can, jointly, complement each other to provide more information about song history than cells with identical context preferences (Figure 4a and Figure 3 ROIs 21, 45, 50, Figure 2d). In our dataset of sparsely labelled neurons, ROIs were rarely active in the same phrase. Figure 4d, shows four ROIs that were jointly active during a single phrase type (pink in Figure 4d). One ROI’s activity was specific to a single context (Figure 4d ROI 10. c.f. as in ^15^) and the other three were active in multiple contexts (Figure 4d, ROIs 24, 29, 31). The phrase during which these ROIs were recorded precedes a complex transition and the type of the preceding phrase poorly predicts the transition outcome (right bar in Figure 4e, 0.08 out of 1 bit, bootstrapped mutual information estimate, methods). However, looking at multiple ROIs together we found that the network holds significantly more information about the past and future phrase types (Figure 4e, 0.42, 0.33 bit, Bootstrapped z-test p<1e-6). This increased predictive power suggests that population responses can contain the long-range information required in the complex transition. Furthermore, in this example the network holds significantly more information about the past than the future (Figure 4e, bootstrapped z-test, p<1e-4, Supplementary Figure 4 - 2), suggesting that information is lost during the complex transition, as also demonstrated by the bias in encoding past contexts by individual ROIs (Figure 2e).

Taken together, these findings demonstrate that neural activity in canary HVC carries long range song context information. These activity patterns can be described as “hidden states” since they relate primarily to past or future song elements and do not alter the syllable class being sung (c.f. Supplementary Figure 1 - 7 illustrating this class distinction for the transition in Figure 1). These patterns of activity contain the information necessary to drive complex, context-dependent phrase transitions.

## Discussion

Motor sequences with long-range order dependencies are common in choreographed behaviors, like language, but the neural mechanisms are largely unknown.

Several landmark experiments demonstrate that HVC projection neuron activity in zebra finches is highly stereotyped during the production of syllables^17,38^. Hidden premotor states^6^ driving the same syllable in different contexts provide tight neural-motor correlations in the moment to moment execution of song syllables while also supporting sequence-dependent transitions at the level of phrases. Here we validate four key predictions of this many-to-one hypothesis, and show that HVC contains the information necessary to follow long range syntax rules; consecutively-active ROIs reporting the identity of phrases up to four steps in the past, ROIs that predict phrase types two steps into the future, concentration of song-context-correlates in complex transitions, and ROIs selective to multiple contexts. These observations resemble the many-to-one relation between neural activity and behavior states^6,29,33^ proposed in some models to relay information across time and, expanding on observations in other species^7–14,39^, support syntactic rules. Our findings also expand on a prior study in Bengalese finches^15^, that showed neural correlations to song-sequence up to 100ms apart, and demonstrate hidden states related to long-range memory-guided syntax rules extending over several seconds.

However, there are also clues that HVC does not contain all the information required to select a phrase transition – since more neurons correlate to the sequence’s past than to its future it is possible that sequence information in HVC is lost, perhaps supporting stochastic transitions. The source of residual stochasticity in HVC could be intrinsic to the dynamics of HVC – for example the “noise” terms commonly added in sequence generating models^24,40,41^ or may enter downstream as well-documented noise sources in the basal ganglia also converge on pre-motor cortical areas downstream of HVC. These early findings require further investigation of the neural dynamics during flexible transitions and may provide a tractable model for studying stochastic cognitive functions – mechanisms in working memory and sensory-motor integration that remain extremely challenging to quantify in most spontaneous behaviors in mammals.

Finally, it is worth noting that recent dramatic progress in speech recognition algorithms have employed recurrent neural networks with several architectures designed to capture sequence dependencies with hidden states. Examples include LSTM^42^, hierarchal time scales^43^, hidden memory relations^44^, and attention networks^45^. It is possible that machine learning models will help to interpret the complex dynamics of HVC, and help inform new models of many to one, history dependent mappings between brain state and behavior^29^.

## Supporting information

Supplemental analysis 1

Supplemental analysis 2

## Supplementary Figures and Legends

**Supplementary Figure 1 - 1.**
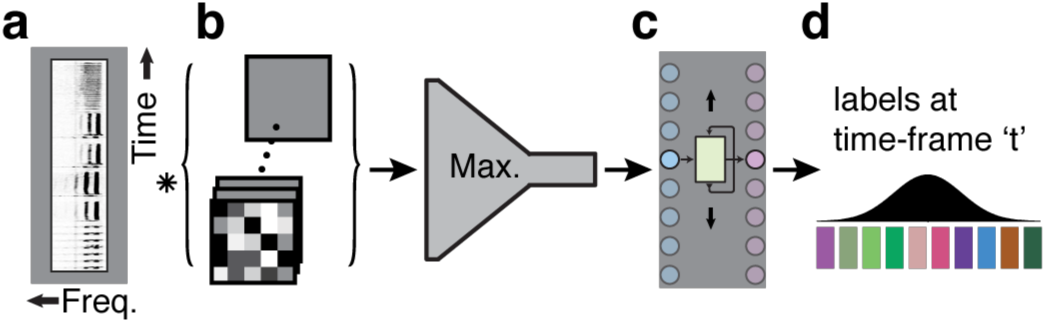
Architecture of syllable segmentation and annotation machine learning algorithm. **a.** A spectrogram as a 2D input matrix is fed to the algorithm in segments of 1 second. **b.** Convolutional and max-pooling layers allow learning local spectral and temporal filters. **c.** Bidirectional recurrent Long-Short-Term-Memory (LSTM) layer allows learning temporal sequencing features. **d.** Projection onto syllable classes assigns a probability for each 2.7 millisecond time bin and syllable.

**Supplementary Figure 1 - 2.**
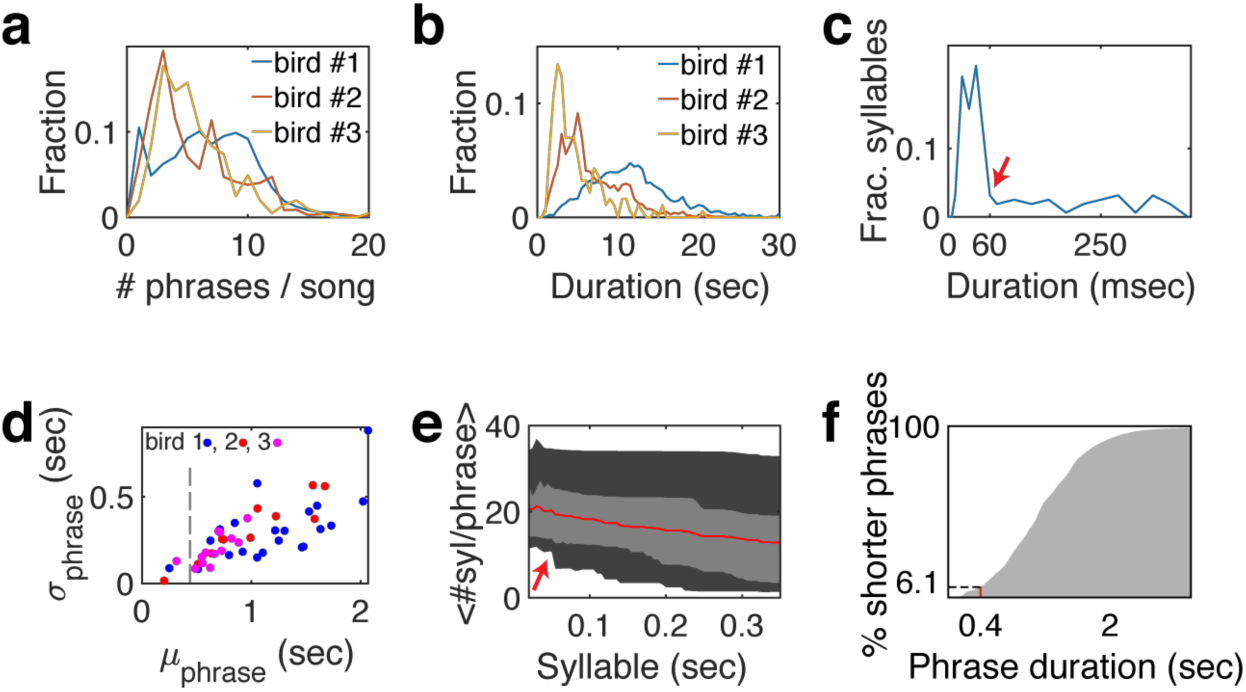
Syllable and phrase statistics. **a.** Histogram of the number of phrases per song for 3 birds used in this study. **b.** Histogram of song durations for 3 birds. **c.** Histogram of mean syllable durations, 85 syllable classes from 3 birds. Red arrow marks the duration, below which all trill types have more than 10 repetitions on average. **d.** Relation between syllable classes’ duration mean (x-axis) and standard deviation (y-axis). Syllables classes (dots) of 3 birds are colored by the bird number. Dashed line marks 450 msec, an upper limit for the decay time constant of GCaMP6f. **e.** Range of mean number of syllables per phrase (y-axis) for all syllable types with mean duration shorter than the x-axis value. Red line is the median, light gray marks the 25%, 75% quantiles and dark gray mark the 5%, 95% quantile. The red arrow matches the arrow in panel c. **f.** Cumulative histogram of trill phrase durations.

**Supplementary Figure 1 - 3.**
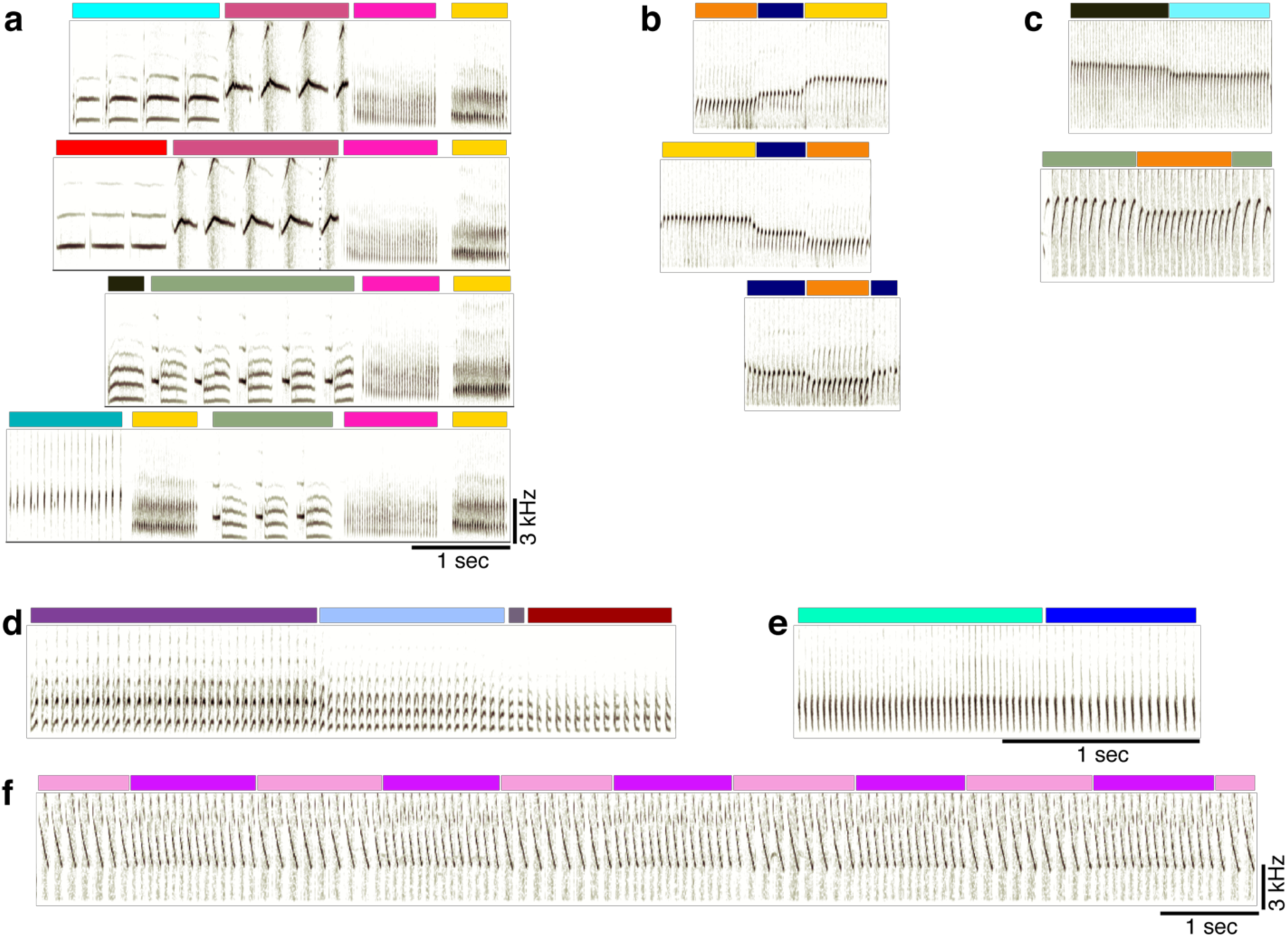
More examples of canary song phrase sequences. **a.** Additional spectrograms of phrase sequences (colors above the spectrograms indicate phrase identity), leading to a repeating pair of phrases (pink and yellow). **b.** Examples of flexible phrase sequencing comprised of pitch changes (from bird #3). **c.** Examples of phrase transitions with a pitch change from bird #2. **d-f.** Phrase sequences showing changes in spectral and temporal parameters. d, bird #1, changes from up sweep (purple) to down sweep (dark red) through intermediate phrases of intermediate acoustic structure. e, bird #1, a change in inter-syllable gaps. f, from bird #2, changes in pitch sweep rate.

**Supplementary Figure 1 - 4.**
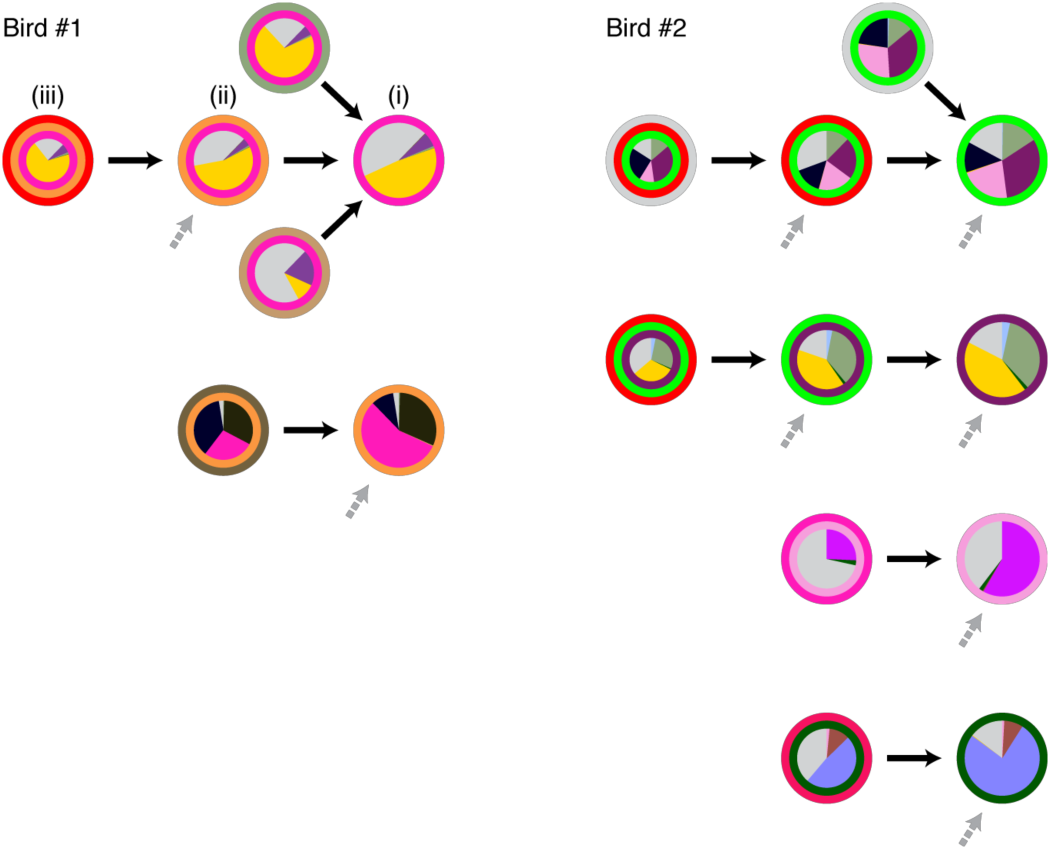
All complex phrase transitions with ≥2^nd^ order dependence on song history context (for birds #1, #2). For each phrase type that precedes a complex transition, the context dependence is visualized by a graph called a Probabilistic Suffix Tree (methods). Transition outcome probabilities are marked by pies at the center of each node. The song context—phrase sequence—that leads to the transition, is marked by concentric circles, the inner most being the phrase type preceding the transition. Nodes are connected to indicate the sequences in which they are added in the search for longer Markov chains that describe context dependence (e.g. i-iii for 1^st^ to 3^rd^ order Markov chains). Grey arrows indicate additional incoming links that are not shown for simplicity.

**Supplementary Figure 1 - 5.**
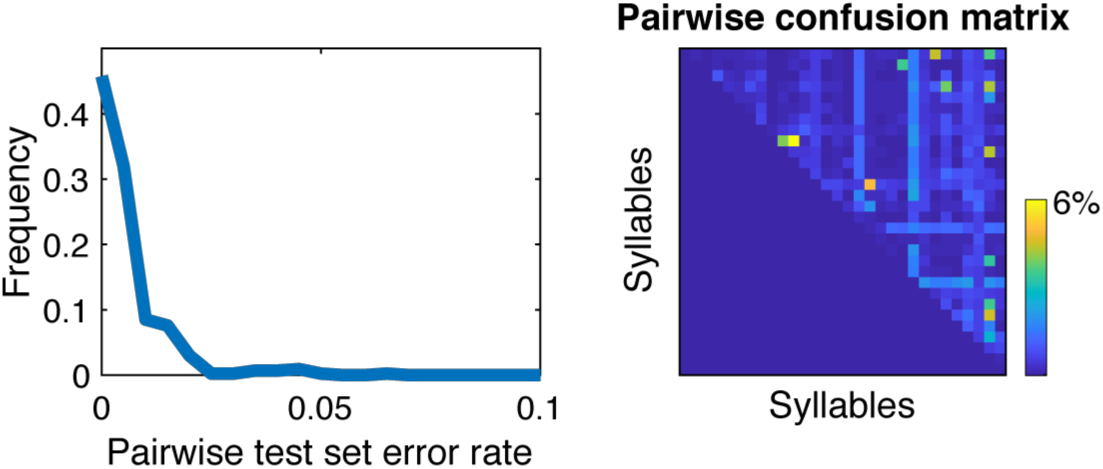
The syllables’ acoustic variability cannot lead to phrase sequence errors. A support vector machine (SVM) classifier was used to assess the pairwise confusion between all syllables classes of bird #1 (methods). The test set confusion matrix (right) and its histogram (left) show that in rare cases the error exceeded 1% and at most reached 6%. Since the higher values occurred only in phrases with 10s of syllables this metric guarantees that most of the syllables in every phrase cannot be confused as belonging to another syllable class. Accordingly, the possibility for making a mistake in identifying a phrase type is negligible.

**Supplementary Figure 1 - 6.**
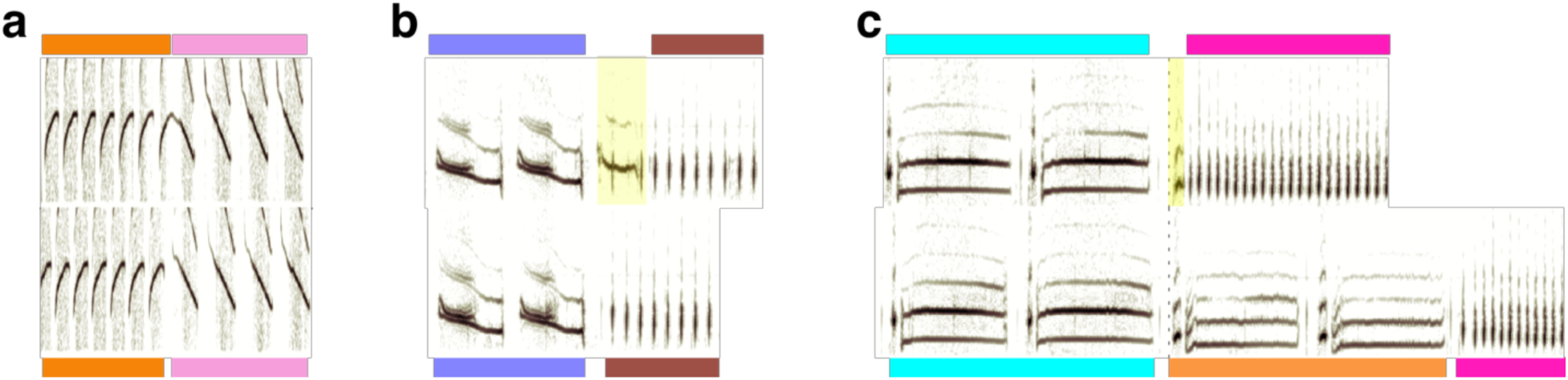
Examples of rare inter-phrase gaps and aberrant syllables. **a.** Top and bottom sonograms compare the same phrase transitions where the inter-phrase gap varies. **b**, **c.** The top sonogram includes a rare vocalization in the beginning of the 2^nd^ phrase (highlighted) that, in panel c, resemble the onset of an orange phrase type.

**Supplementary Figure 1 - 7.**
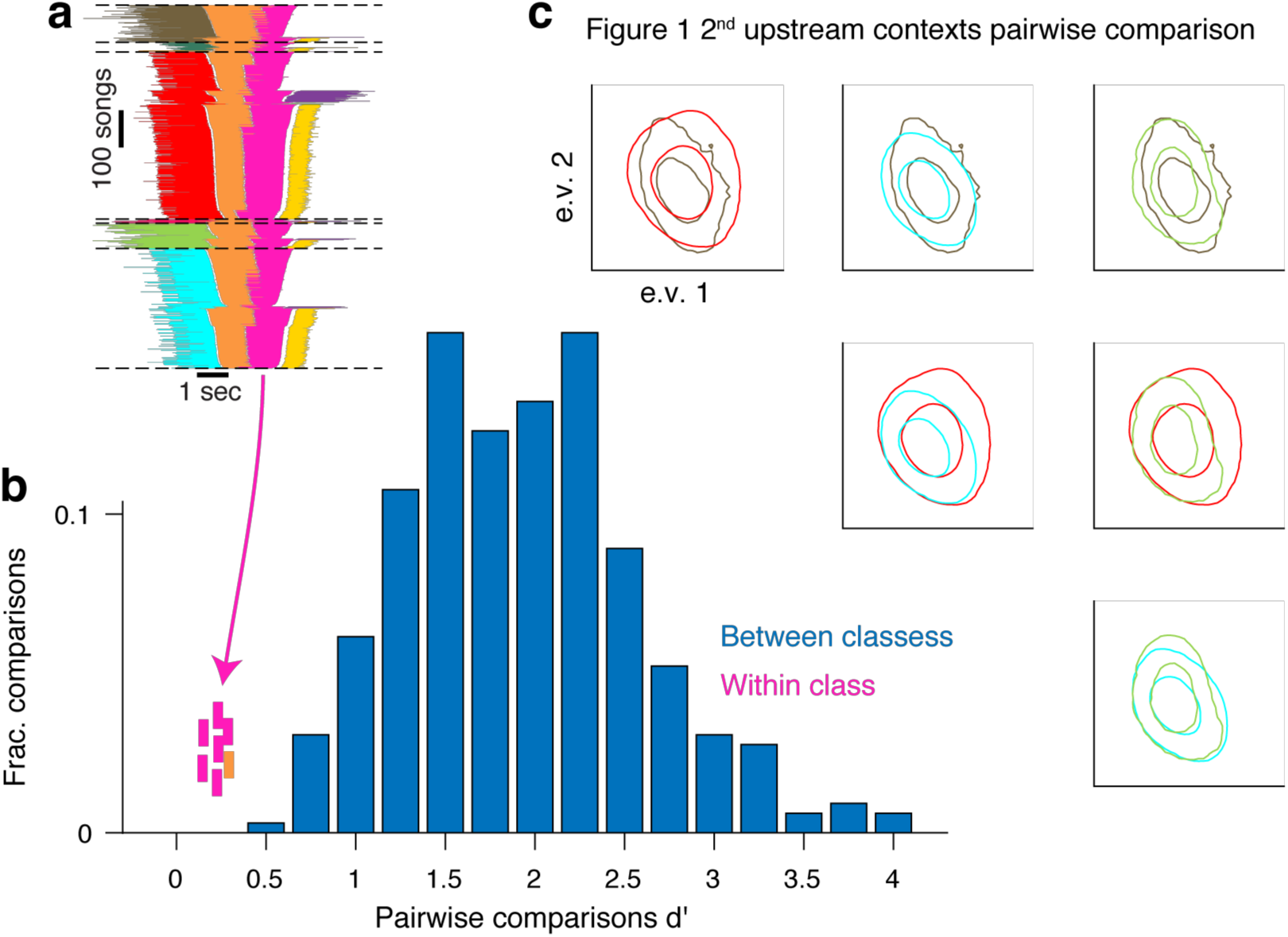
An example in which context-dependence of syllable acoustics prior to complex transitions is too small for clear distinction. **a.** Repeats main figure 1b. A summary of all phrase sequences that contain a common transition reveals that the choice of what to sing after the pink phrase depends on the phrases that were produced earlier. Lines represent phrase identity and duration. Song sequences are stacked (vertical axis) sorted by the identity of the 1st phrase, the last phrase and then the center phrases’ duration. **b.** The discriminability (d’, x-axis) measures the acoustic distance between pairs of syllable classes in units of the within-class standard deviation (methods). Bars show the histogram across all pairs of syllables identified by human observers (methods) corresponding to about 99% or larger identification success (in Supp. Figure 1-5). The pink ticks mark the d’ values for 6 within-class comparison of the main 4 contexts in panel a. The orange tick marks the d’ another context comparison in a different syllable that precedes a complex transition for this bird. **c.** The pairwise comparison of distributions matching the pink ticks in panel b. Each inset shows overlays of two distributions marked by contours at the 0.1 and 0.5 values of the peak and colored by the context in panel a. The distributions are projected onto the 2 leading principle components of the acoustic features (methods). While some of these distributions are statistically distinct they only allow for ~70% context identification success in the most distinct case.

**Supplementary Figure 2 - 1.**
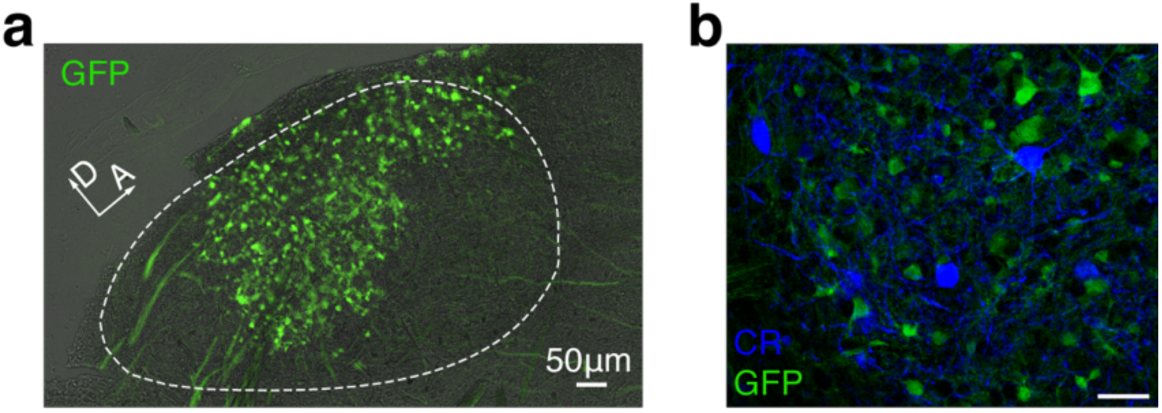
Calcium indicator is expressed exclusively in HVC excitatory neurons. **a.** Sagittal slice of HVC showing GCaMP expressing projection neurons. **b.** We observed no overlap between transduced GCaMP6f-expressing neurons, and neurons stained for the inhibitory neurons markers calretinin, calbindin, and parvalbumin (CR stain shown).

**Supplementary Figure 2 - 2.**
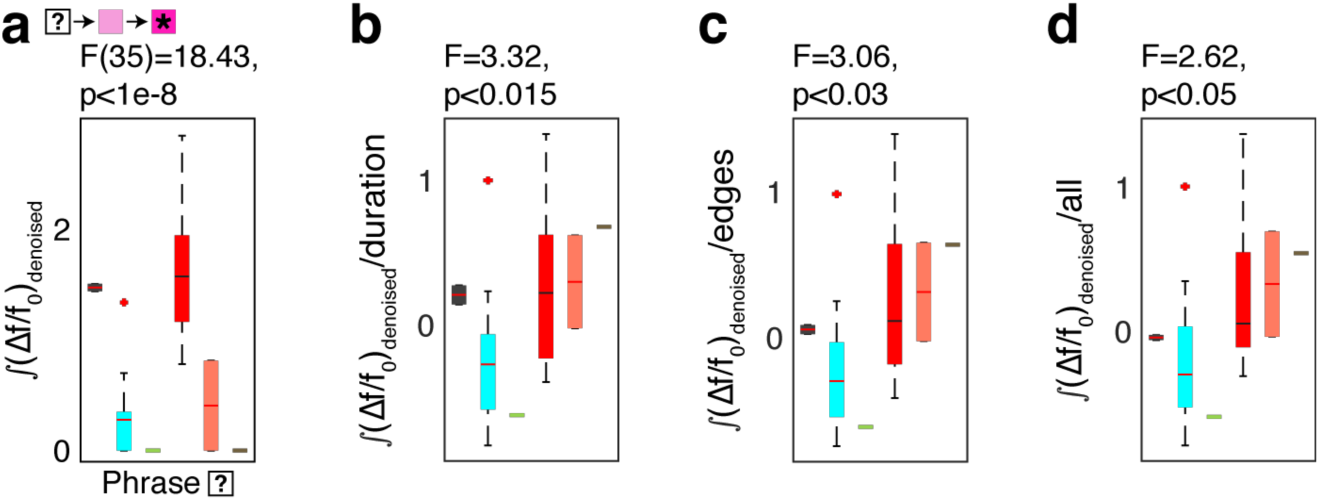
Statistics of one example sequence-correlated neuron. **a.** Quantification of the 2^nd^ order context correlation for the neuron in Figure 2d using 1-way ANOVA (F,p), to test the effect of contexts (x-axis, 2^nd^ preceding phrase type) on the signal integral (y-axis, ∫ (*Δf*/*f*_0_)_*denoised*_), that occurs during the target phrase (marked by ★). **b-d.** The sequence correlation in panel (a) remains significant after accounting for confounding variables. ANOVA tests are carried out using the residuals from the signal integral during the target phrase after removing the cumulative linear dependence on: **b.** The duration of the target phrase. **c.** The relative timing of onset and offset edges of two fixed phrases. **d.** Also including the absolute onset time of the target phrase in each rendition. Colors in this figure correspond to phrases represented in Figure 2d.

**Supplementary Figure 2 - 3.**
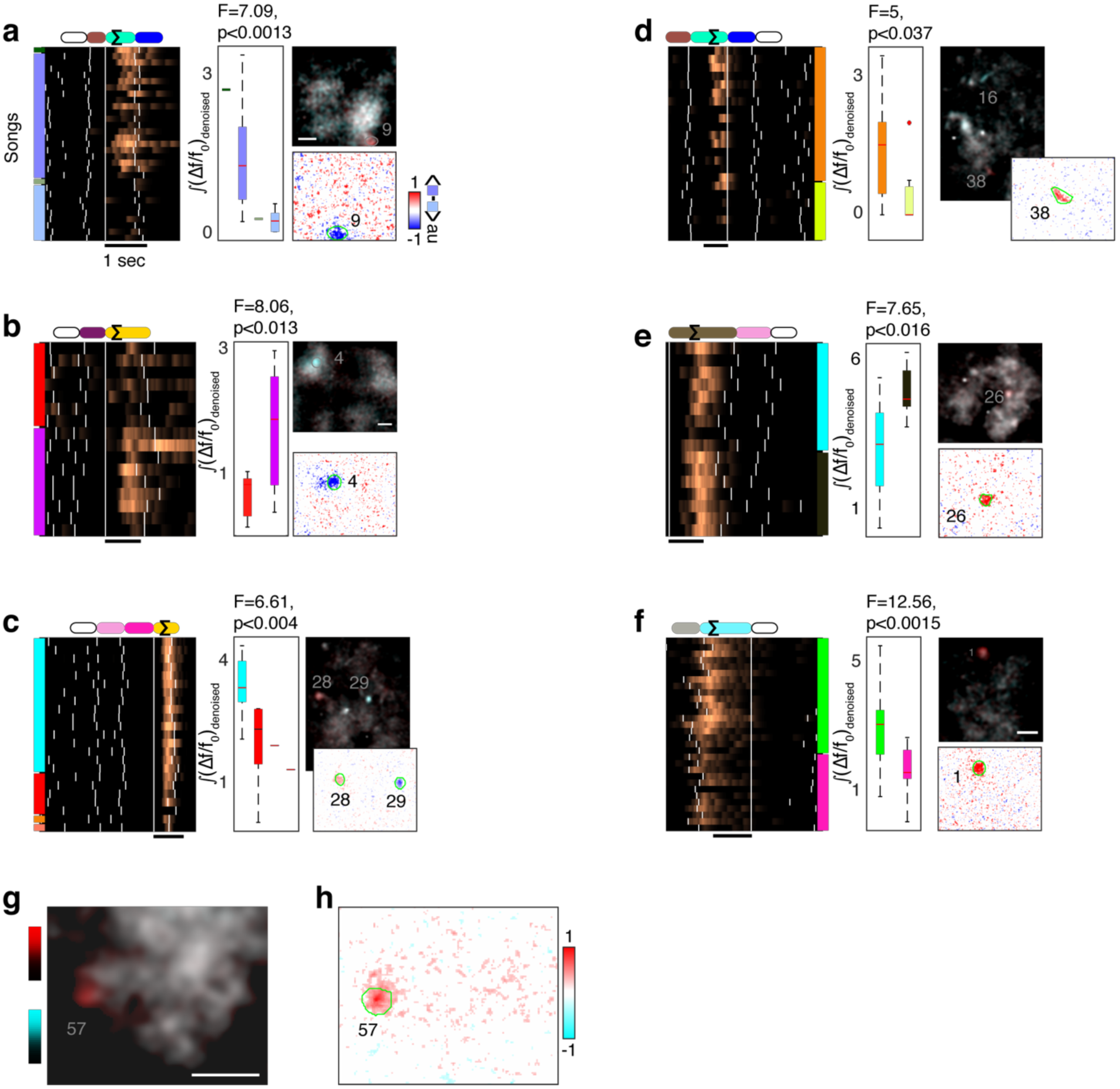
Additional examples of phrase-sequence-correlated ROIs from 3 birds. ROI activity during a target phrase (marked by Σ) that is strongly related to non-adjacent phrase identities (empty ovals in color coded phrase sequence above the raster). Neural activity is aligned to the onset of the current phrase. Songs are arranged by the phrase sequence context (left or right color patches for past and future phrase types). White ticks indicate phrase onsets. Box plots show the 1-way ANOVA (F,p), used to test the effect of contexts (x-axis, phrase type in the empty oval) on the signal integral (y-axis, ∫ (*Δf*/*f*_0_)_*denoised*_), that occurs during the target phrase. Additionally, the context-dependent ROIs are visualized by comparing maximum projection images in the contrasted contexts (methods). In the top insets in panels a-f and in panel g, the projection images are overlaid in orthogonal colors (red and cyan) to reveal context-dependent regions in color. In the bottom insets in panels a-f and in panel h, the maximum projection images are subtracted. **a**,**b.** Similar to main Figure 2d, (*Δf*/*f*_0_)_*denoised*_ from ROIs with 2^nd^ order upstream sequence (color coded) from two more birds. **c.** 3^rd^ order upstream relation. **d**,**e.** 2^nd^ order downstream relations. **f.** 1^st^ order downstream relation from another bird. **g**. Average maximum fluorescence images during the pink phrase in Figure 2d, compare the two most common contexts in orthogonal colors (red and cyan). Scale bar is 50*μ*m. **h.** The difference of the overlaid images in panel g. ROI 57 outlined in green.

**Supplementary Figure 2 - 4.**
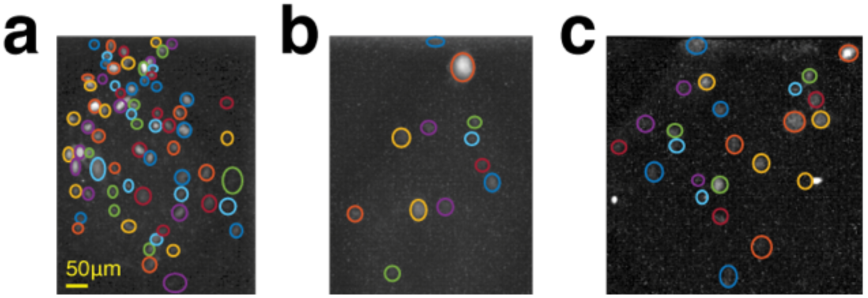
Example of daily ROI annotation in 3 birds. Colored circles mark different ROIs, manually annotated on maximum fluorescence projection images an exemplary day (see methods). Panel **a-c** are for birds 1-3.

**Supplementary Figure 2 - 5.**
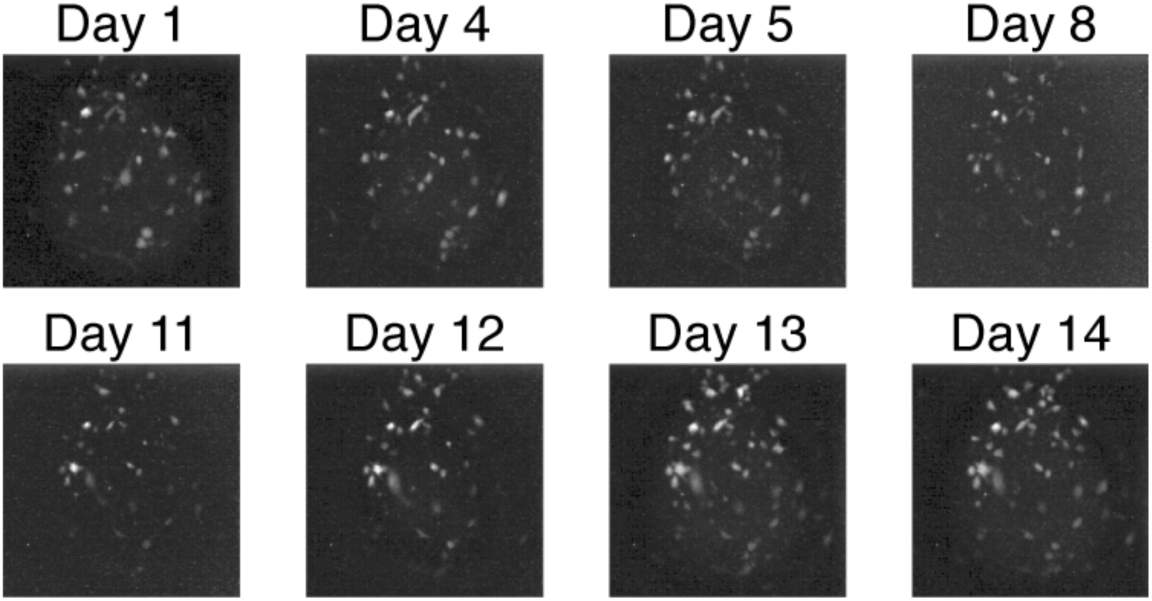
Maximum projection images over 2 weeks (bird #1). Maximum fluorescence images (methods) revealing the fluorescence sources including sparsely active cells in the imaging window across multiple days.

**Supplementary Figure 2 - 6.**
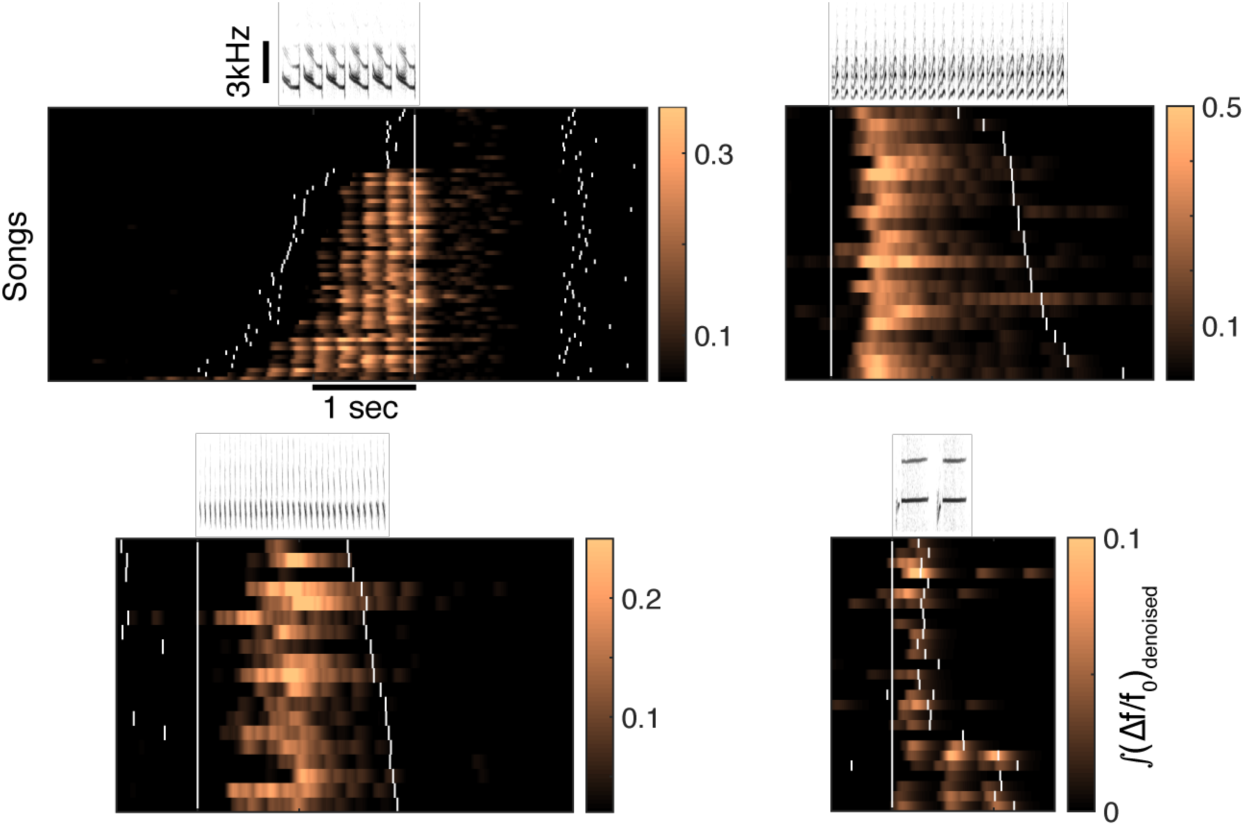
Examples of syllable-locked and phrase-locked cells. Panels show sonogram samples (3kHz frequency scale bar) on top of rasters from 4 ROIs from 3 birds. White ticks indicate phrase onsets. The timescale of the fluorescent calcium indicator is able to resolve individual syllables for the longer syllables.

**Supplementary Figure 2 - 7.**
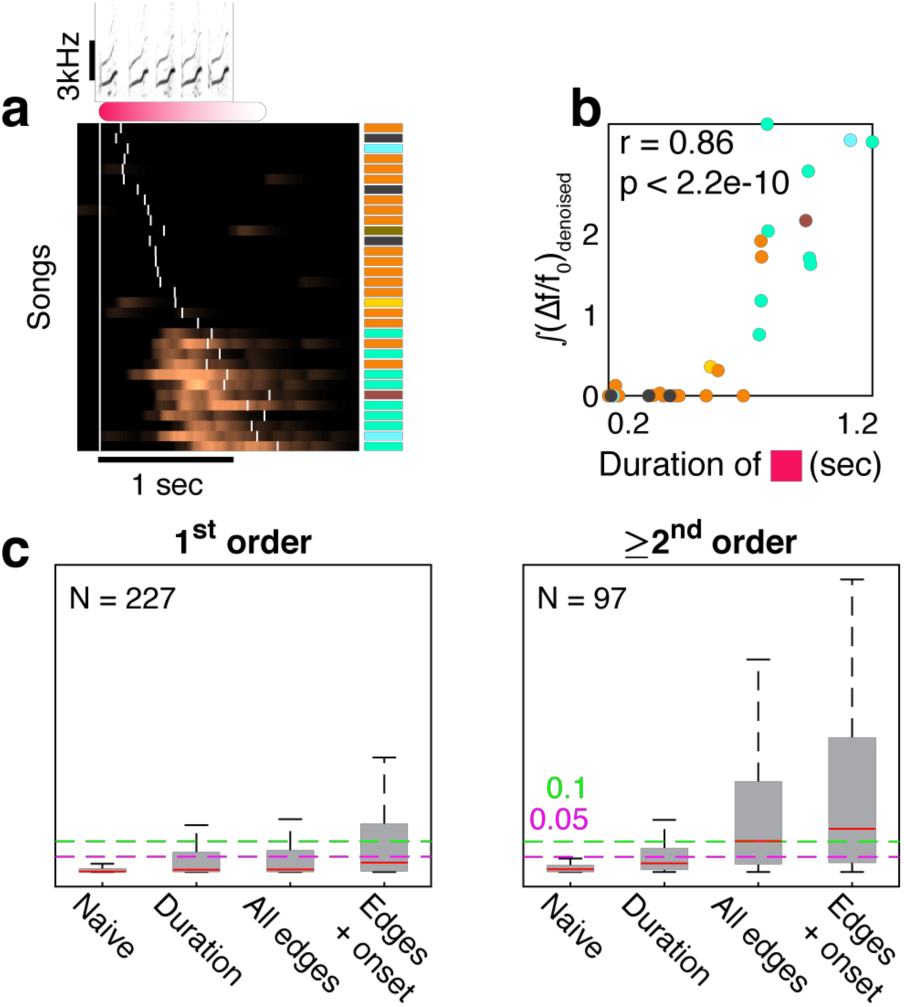
Phrases’ durations and onset times also correlate to the phrase sequence, but cannot fully account for HVC activity. a. (*Δf*/*f*_0_)_*denoised*_ traces (ROI #18, bird #3) during a single phrase type (red) arranged by its duration. Colored patches mark the identity of the final phrase in the sequence. **b.** Correlation between the signal integral and the red phrase duration (markers are colored by the identity of the following phrase, matching the barcode in panel a, Pearson r,p values). **c.** Distributions of 1-way ANOVA p-values (y-axis) relating phrase identity and signal integral for adjacent phrases (1^st^ order transitions, left) and non-adjacent phrases (≥2^nd^ order, right). Tests are also done on residuals of signal integrals, after discounting the following variables: variance explained by the target phrase duration, the timing of all phrase edges in the test sequence, and the time-in-song (x-axis, effects accumulated left to right by multivariate linear regression, see methods). Colored, dashed lines mark 0.05 and 0.1 p-values.

**Supplementary Figure 2 - 8.**
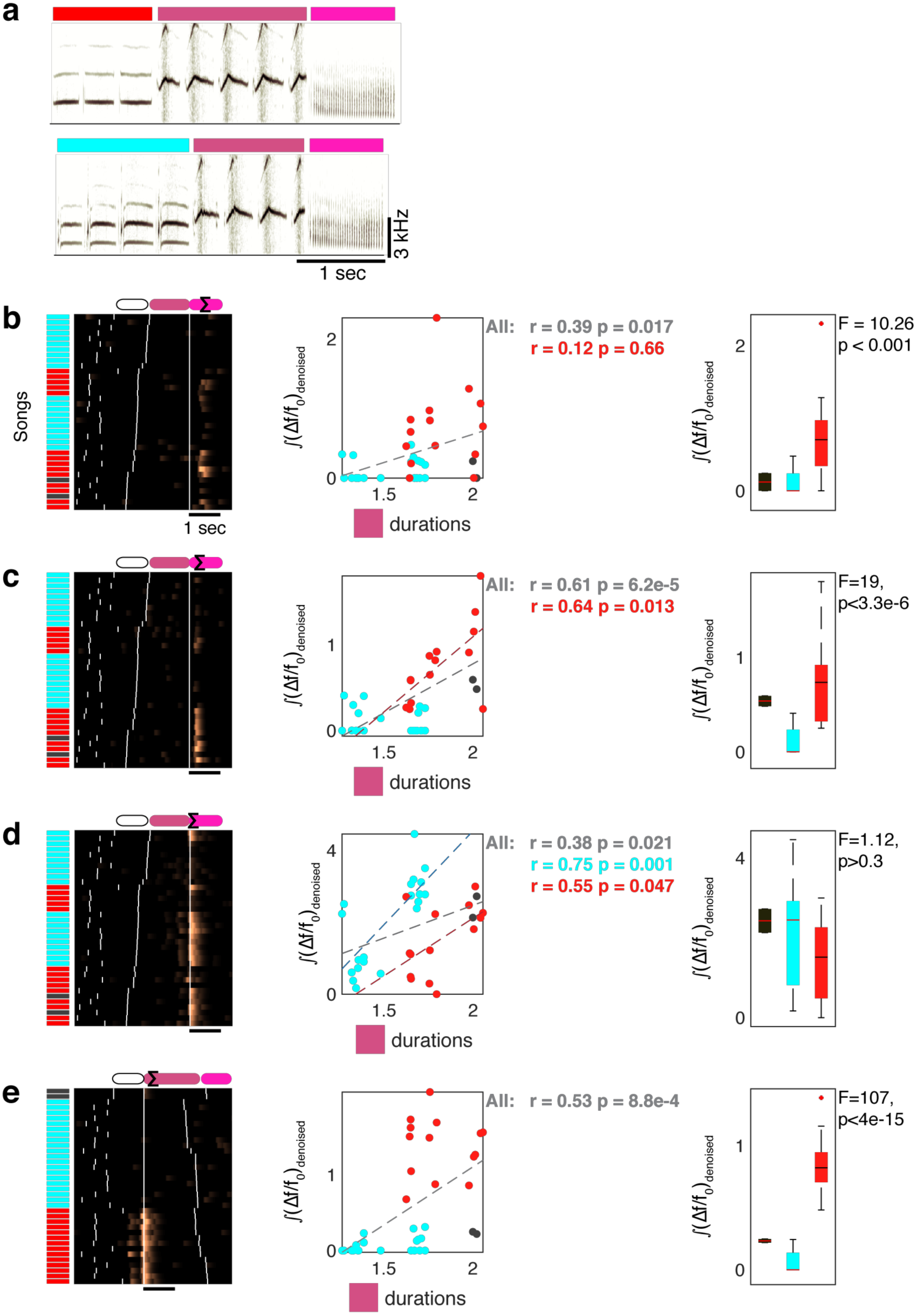
Additional examples of the relationship between neural activity, sequence and duration in 4 ROIs recorded in the same day. **a.** Sonograms show two main phrase sequences. **b-e.** Neural activity of 4 ROIs are aligned to the middle phrase (panels b-d to its offset, panel e to its onset). Songs (y-axis) are ordered by the duration of the middle phrase (b-d) or the identity of the first phrase (e, color coded on left side) and show a variety of properties that yield signal correlation to the duration of the middle phrase (scatter plots, dashed lines indicate significant correlations. Pearson r,p values and markers are colored by the phrase type they represent). Box plots show 1-way ANOVA (F,p) between the signal integral (Σ) in each song and the label of the 1^st^ phrase (color). **b.** In this cell, the signal correlation with the intermediate phrase duration is completely entangled with the signal’s sequence preference and does not apply in separate preceding contexts (red, p > 0.5). **c.** In this cell, the signal correlation with the intermediate phrase duration is influenced by the signal’s sequence preference but also exists in the preferred sequence context separately (red). **d.** For this cell, the signal’s duration correlation is observed within each single preceding context separately, but the correlation reduces across all songs. **e.** Similar to panel a, but the signal is in the 2^nd^ phrase, not the 3^rd^.

**Supplementary Figure 2 - 9.**
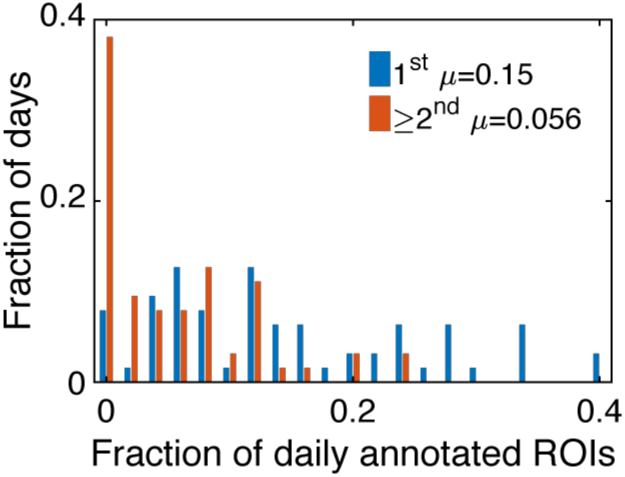
Histogram of fractions of daily annotated ROIs showing sequence correlation in all 3 birds. Colors separate 1^st^ order relations from all the rest. Each ROI can be counted as having both 1^st^ and ≥2^nd^ sequence correlations. A ROI that has sequence correlation of the same order in more than one context is counted only once. This estimate includes all ROIs, including sparsely active ones (c.f. Supplementary Figure 2 - 5).

**Supplementary Figure 3 - 1.**
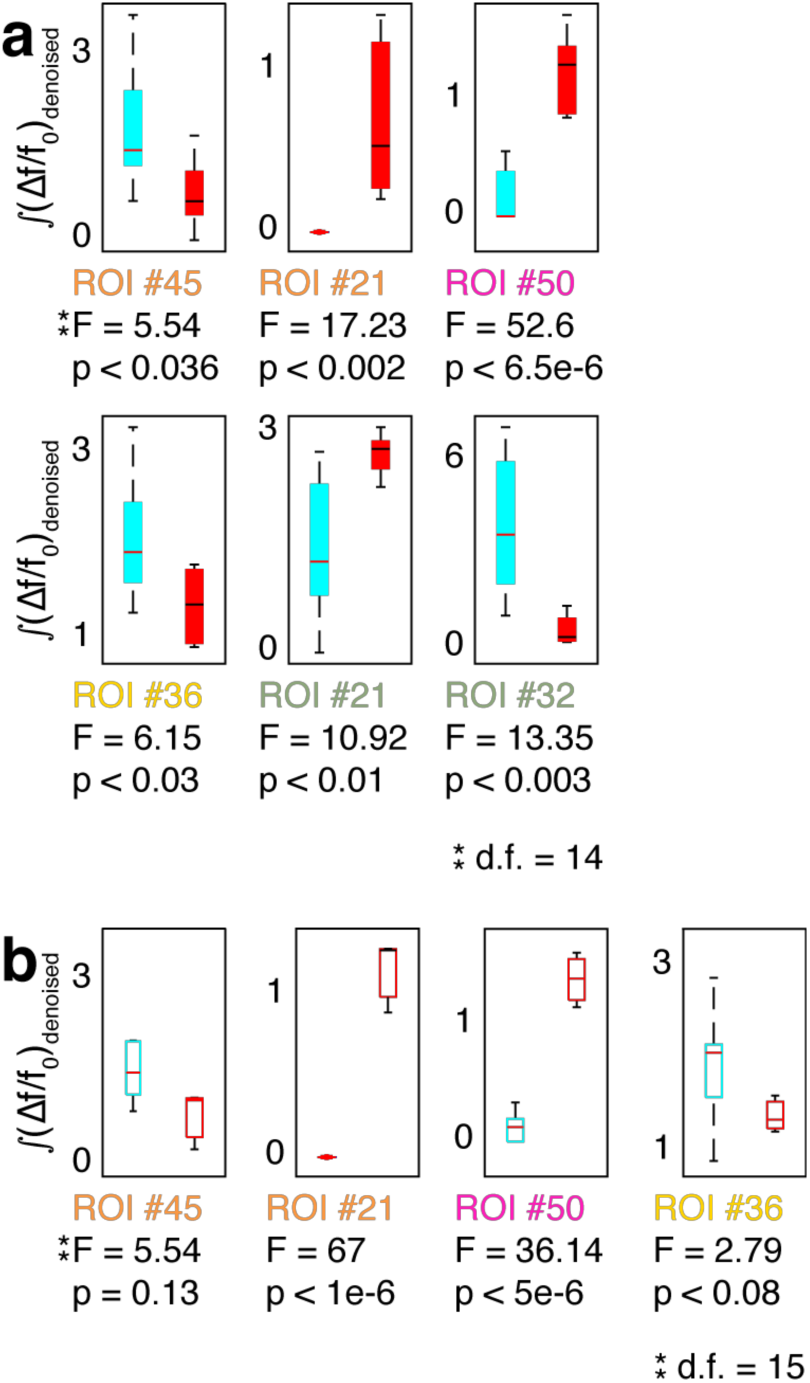
Statistical tests for ROIs in Figure 3. **a.** Distribution of signal integrals (y-axis) for ROIs in Figure 3a. (Text label is color coded by phrase type in sub-panels i-iv). F-numbers and p-values for 1-way ANOVA relating history (x-axis) and signal (y-axis). **b.** ROIs in panel (a) retain their song-context bias also for songs that happen to terminate at end of the third phrase rather than continue. Box plots repeat the ANOVA tests in panel (a) for songs in which the last phrase is replaced by end-of-song.

**Supplementary Figure 3 - 2.**
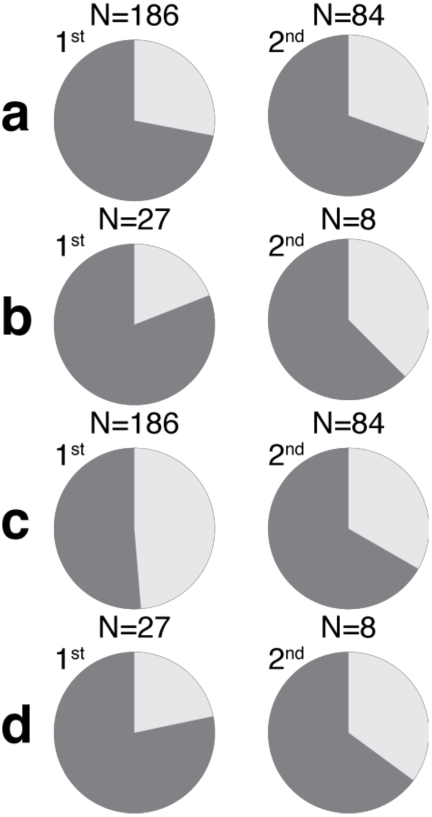
ROIs with significant context-dependent correlations are more frequently found in complex transitions - points in the behavior that depend on long range context– data from 2 birds. Dark grey indicates the fraction of correlations occurring in complex behavioral transitions. **a**,**b.** the data in Figure 3c separated to the two birds. **c**,**d.** Normalizes the distributions in panels a,b by the frequency of each phrase type.

**Supplementary Figure 3 - 3.**
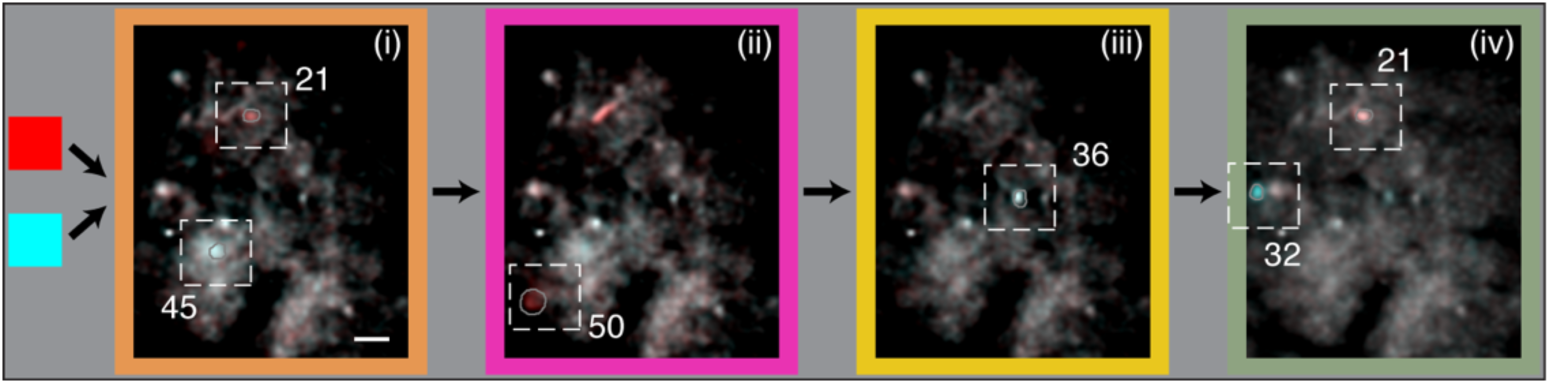
Maximum fluorescence images replicate the context sensitive ROIs in de-noised video data. Figure 3a showed maximum projection images, calculated with de-noised videos (methods). The algorithm, CNMF-E^46^, involves estimating the source ROI shapes, de-convolving spike times as well as estimating the background noise. Here, recreating the maximum projection images with the original fluorescence videos shows the background as well but the preceding-context-sensitive neurons remain the same. Namely, the same ROI footprints annotated in panels i-iv show the color bias (cyan or red) that indicates coding of the past phrase with the same color.

**Supplementary Figure 3 - 4.**
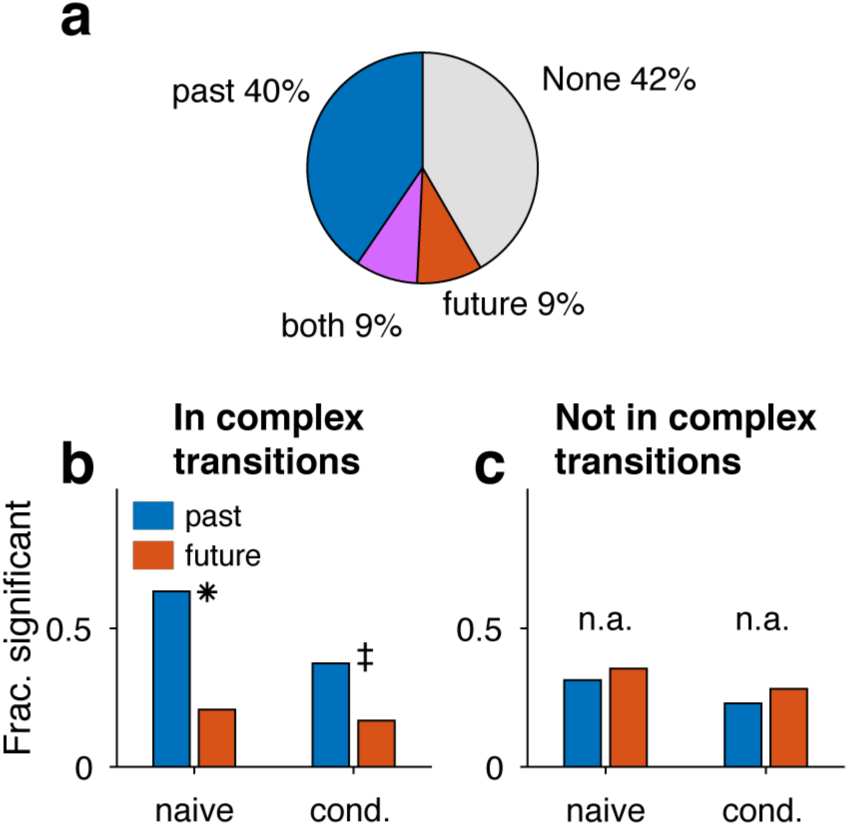
Sequence-correlated ROIs in complex transitions are biased to represent past contexts. **a.** In sequence-correlated ROIs, multi-way ANOVA is used to separate the effect of the preceding and following phrase types on the signal (methods). Pie shows the percent of sequence-correlated ROIs significantly influenced by the past, future, or both phrase identities. **b.** Restricting analysis to complex transitions, more ROIs correlated to the preceding phrase type (blue) than to following (red). This is true in both Naive signal values (left) and after removing dependencies on phrase durations and time-in-song (right). (binomial z-test: *: p < 1e-13, ‡: p < 0.0001). **c.** Restricting to phrase types not in complex transitions reveals More ROIs correlated with the future phrase type but the difference is not significant (binomial z-test, p > 0.2)

**Supplementary Figure 4 - 1.**
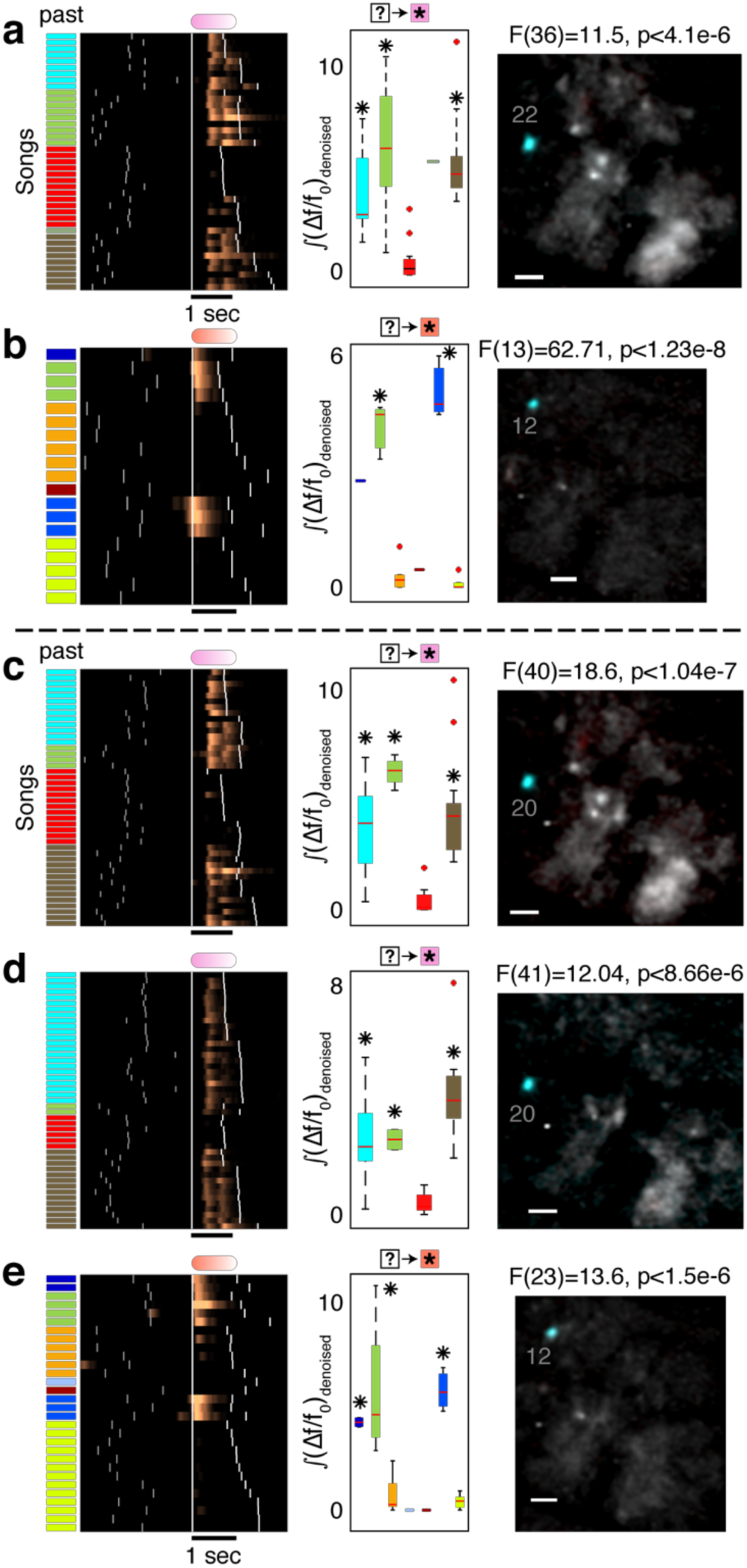
ROIs in Figure 4 reflect several preceding song contexts. **a**,**b.** Rasters repeat Figure 4a,b. Box plot shows distributions of (*Δf*/*f*_0_)_*denoised*_ integrals (y-axis, summation in the phrase marked by ★) for various song contexts (x-axis). F-number and p-value show the significance of separation by song context (1-way ANOVA) and * marks contexts that lead to larger mean activity compared to another context (Tukey’s multiple comparisons, p<0.001). Average maximum projection images (methods) during the aligned phrase compare the song contexts that lead to significantly higher activity to the other contexts in orthogonal colors (cyan and red for high and low activity). Bar is 50*μ*m. **c-e.** Neurons with similar context preference like the examples in Figure 4a,b in adjacent days.

**Supplementary Figure 4 - 2.**
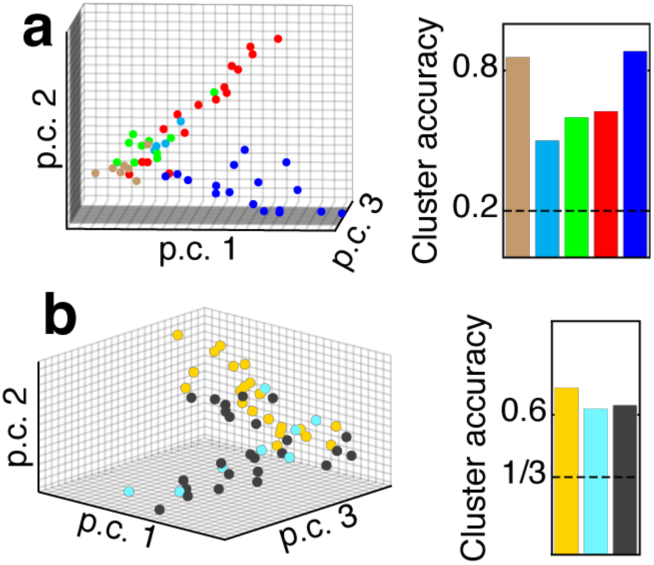
The 4-ROI network states in Figure 4d form more separable clusters when labeled by the past context compared to the future context. **a.** Signal integrals from the 4 ROIs in Figure 4d are plotted for each song (dots) on the 3 most informative principle components. Dots are colored by the identity of the preceding phrase. Clustering accuracy measures the ‘leave-one-out’ label prediction for each preceding phrase (true positive), calculated by assigning each dot to the nearest centroid (L_2_). Dashed line marks chance level. **b.** Similar to panel a but for the 1^st^ following phrase.

## Supplementary videos

### Supplementary videos 1-7

Video frames show stacks of confocal microscopy section (3µm thick) that were used to test the specificity of GCaMP expression to excitatory neurons (methods). In all movies, GCaMP is stained in green and the inhibitory neuron markers (CB,CR,PV annotated in the file names) are stained in blue.

### Supplementary video 8

The results of the CNMFE^46^ algorithm that was used to de-noise the fluorescence videos to visualize context dependent neurons in Figure 3a.

## Acknowledgements

The authors would like to thank Jeff Markowitz, Ian Davison, and Jeff Gavornik for useful discussions and comments to this manuscript.

## Materials and Methods

### Ethics declaration

All procedures were approved by the Institutional Animal Care and Use Committee of Boston University (protocol numbers 14-028 and 14-029). Imaging data were collected from *n* = 3 adult male canaries. Birds were individually housed for the entire duration of the experiment and kept on a light–dark cycle matching the daylight cycle in Boston (42.3601 N). The birds were not used in any other experiments.

### Data availability

The datasets are available from the corresponding author on request.

### Code availability

All custom-made code in this manuscript is publicly available in Github repositories. URLs are provided in the relevant methods descriptions

### Surgical procedures

#### Anesthesia and analgesia

Prior to anesthetizing the birds, they were injected with meloxicam (intramuscular, 0.5mg/kg) and deprived of food and water for a minimum of 30 minutes. Birds were anesthetized with 4% isoflurane and maintained at 1–2% for the course of the surgery. Prior to skin incision, bupivacaine (4 mg/kg in sterile saline) injected subcutaneously (volume 0.1-0.2 mL). Meloxicam was also administered for 3 days after surgery.

#### Stereotactic coordinates

The head was held in a previously described, small animal stereotactic instrument^47^. To increase anatomical accuracy and ease of access, we deviated from the published atlas coordinates^47^ and adapted the head angle reference to a commonly used forehead landmark parallel to the horizontal plane. The outer bone leaflet above the prominent *λ* sinus was removed and the medial (positive = right) and anterior (positive) coordinates are measured from that point. The depth is measured from the brain’s dura surface. The following coordinates were used (multiple values indicate multiple injections):

|  | HVC | RA | Area X |
| --- | --- | --- | --- |
| Head Angle | +65° | +80° | +20° |
| Medial (mm) | -2.5 | -2.5 | -1.27, -1.3 |
| Anterior (mm) | 0.12 | -1.2 | 5.65, 5.8 |
| Depth (mm) | 0.15-0.7 | 1.9-3 | 2.65-2.95 |

#### HVC demarcation and head anchoring

To target HVC, 50-100nL of the DiI retrograde lipophilic tracer (5mg/ml solution in dimethylformamide, DMF) was injected into the left area X. The outer bone leaflet was removed above area X with a dental drill. The inner bone leaflet was thinned and removed with an ophthalmic scalpel, exposing a hole of ~300 µm diameter. The left area X was injected using a Drummond Nanoject II (Drummond pipette, 23nl/sec, pulses of 2.3nl). In the same surgery, a head anchoring structure was created by curing dental acrylic (Flow-It ALC, Pentron) above the exposed skull and through ~100µm holes in the outer bone leaflet.

#### Virus injection and lens implants

A lentivirus that was developed for previous work in zebra finches (containing the vector pHAGE-RSV-GCaMP6f; Addgene plasmid #80315) was also used in canaries^48^. The outer skull leaflet above HVC was removed with a dental drill. The inner bone leaflet was thinned and removed with an ophthalmic scalpel, exposing ~1.5-2mm diameter area of the dura. The DiI demarcation of HVC was used to select an area for imaging. The lentivirus was injected in 3-4 locations, at least 0.2mm apart, at a range of depths between 0.5-0.15mm. In total 800-1000nL were injected into the left HVC. After the injection, the dura was removed and the parahippocampus segment above the imaging site was removed with a dura pick and a custom tissue suction nozzle. A relay GRIN lens (Grintech GT-IFRL-100, 0.44 pitch length, 0.47 NA) was immediately positioned on top of the exposed HVC and held in place with Kwik-Sil (WPI). Dental acrylic (Flow-It, Pentron) was used to attach the lens to the head plate and cover the surgery area. The birds were allowed to recover for 1-2 weeks.

#### Hardware

To image calcium activity in HVC projection neurons during singing, we employed custom, lightweight (~1.8 g), commutable, 3D-printed, single-photon head-mounted fluorescent microscopes that simultaneously record audio and video (Figure 2). These microscopes enabled recording hundreds of songs per day, and all songs were recorded from birds longitudinally in their home cage, without requiring adjustment or removal of the microscope during the imaging period. Birds were imaged for less than 30 min total on each imaging day, and LED activation and video acquisition were triggered on song using previously described methods^48^.

#### Microscope design

We used a custom, open-source microscope developed in the lab^48^. A blue LED produces excitation light (470-nm peak, LUXEON Rebel). A drum lens collects the LED emission, which passes through a 4 mm × 4 mm excitation filter, deflects off a dichroic mirror, and enters the imaging pathway via a 0.25 pitch gradient refractive index (GRIN) objective lens. Fluorescence from the sample returns through the objective, the dichroic, an emission filter, and an achromatic doublet lens that focuses the image onto an analog CMOS sensor with 640 × 480 pixels mounted on a PCB that also integrates a microphone. The frame rate of the camera is 30 Hz, and the field of view is approximately 800 μm × 600 μm. The housing is made of 3D-printed material (Formlabs, black resin). A total of 5 electrical wires run out from the camera: one wire each for camera power, ground, audio, NTSC analog video and LED power. These wires run through a custom flex-PCB interconnect (Rigiflex) up to a custom-built active commutator. The NTSC video signal and analog audio are digitized through a USB frame-grabber. Custom software written in the Swift programming language running on the macOS operating system (version 10.10) leverages native AVFoundation frameworks to communicate with the USB frame-grabber and capture the synchronized audio–video stream. Video and audio are written to disk in MPEG-4 container files with video encoded at full resolution using either H.264 or lossless MJPEG Open DML codecs and audio encoded using the AAC codec with a 48-kHz sampling rate. All schematics and code can be found online https://github.com/gardner-lab/FinchScope and https://github.com/gardner-lab/video-capture.

#### Microscope positioning and focusing

Animals were anesthetized and head fixed. The miniaturized microscope was held by a manipulator and positioned above the relay lens. The objective distance above the relay was set such that blood vessels and GCaMP6f expressing cells were in focus. The birds recovered in the recording setup. Within the first couple of weeks, the microscopes were refocused to maximize the number of observable neurons.

#### Histology

DiI was injected into area X as described above. Three days later, ~800nL lentivirus was injected into HVC using the DiI demarcation. In finches, this virus infected predominately projection neurons. In this project we analyzed neurons with sparse activity that do not match the tonic activity of interneurons in HVC. The virus was injected in four sites, at least 0.2mm apart and at two depths (matching the experiment’s procedure). About four weeks later the bird was euthanized (by an intracholemic injection of 0.2mL 10% Euthasol, Virbac, ANADA #200-071, in saline) and perfused by first running saline and then 4% paraformaldehyde via the heart’s left chamber and the contralateral neck vein. The brain was extracted and kept overnight in 4% paraformaldehyde at 4°C.

##### GCaMP6f expression (Supplementary Figure 2 - 1a)

The fixed tissue was sectioned into 70 µm sagittal slices (Vibratome series 1000), placed on microscope slides, and sealed with cover slips and nail polish. Epifluorescence images were taken with Nikon Eclipse Ni-E tabletop microscope.

##### Expression specificity to excitatory neurons (Supplementary Figure 2 - 1b)

The fixed tissue was immersed in 20% and 30% sucrose solutions for two overnights, frozen and sectioned into 30 µm sagittal slices (Cryostat, Leica CM3050S). Following work in zebra finches^49^, the slices were stained for calcium binding interneuron markers Calbindin (1:4000, SWANT), Calretinin (1:15000, SWANT), and Parvalbumin (1:1000, SWANT) by overnight incubation with the primary antibody at 4°C and with a secondary antibody (coupled to Alexa Fluor 647) for 2 hours at room temperature. Slices were mounted on microscope slides, and sealed with cover slips and nail polish. A confocal microscope (Nikon C2si) was used to image GCaMP6f and the interneuron markers in 3µm-thick sections through the tissue. The images were inspected for co-stained cells (e.g. supplementary videos 1-7). The results ruled out any co-expression of GCaMP and Calbindin or Calretinin. We found 2 cells expressing Parvalbumin and GCaMP (supplementary video 5 shows one example, <0.5% of PV stained cells, <0.01% of GCaMP expressing cells), possibly replicating previous observation of PV expression in HVC projection neurons^49^.

### Data collection

#### Song screening

Birds were individually housed in soundproof boxes and recorded for 3-5 days (Audio-Technica AT831B Lavalier Condenser Microphone, M-Audio Octane amplifiers, HDSPe RayDAT sound card and VOS games’ Boom Recorder software on a Mac Pro desktop computer). In-house software was used to detect and save only sound segments that contained vocalizations. These recordings were used to select subjects that are copious singers (>=50 songs per day) and produce at least 10 different syllable types.

#### Video and audio recording

All data used in this manuscript was acquired between late February and early July – a period during which canaries perform their mating season songs. To avoid over exposure of the fluorescent proteins, data collection was done during the morning hours (from sunrise until about 10am) and the daily accumulated LED-on time rarely exceeded 30 minutes. Audio and video data collection was triggered by the onset of song as previously described^48^ with an additional threshold on the spectral entropy that improved detection of song periods dramatically. Data files from the first couple of weeks, a period during which the microscope focusing took place and the birds sang very little, was not used in this manuscript. Additionally, data files from (extremely rare) days in which video files were corrupted because of tethering malfunctions, were not used in this manuscript.

### Data Analysis

#### Video file preprocessing

Software developed in-house was used to load video frames and audio signal to MATLAB (https://github.com/gardner-lab/FinchScope/tree/master/Analysis%20Pipeline/extractmedia) along with the accompanying timestamps. Video frames were interpolated in time and aligned to an average frame rate of 30Hz. Audio samples were aligned and trimmed in sync with the interpolated frame timestamps. To remove out-of-focus bulk fluorescence from the 3-dimensional representation of the video (rows × columns × frames), the background was subtracted from each frame by smoothing it with a 145 pixel-wide circular Gaussian kernel, resulting in 3 dimensional video data, *V x, y, t*.

#### Audio processing

Song syllables were segmented and annotated in a semi-automatic process:

- A set of ~100 songs was manually annotated using a GUI developed in-house (https://github.com/yardencsGitHub/BirdSongBout/tree/master/helpers/GUI). This set was chosen to include all potential syllable types as well as cage noises.
- The manually labeled set was used to train a deep learning algorithm developed in-house (https://github.com/yardencsGitHub/tweetynet).
- The trained algorithm annotated the rest of the data and its results were manually verified and corrected.
- In both the training phase of TweetyNet and the prediction phase for new annotations, data is fed to TweetyNet in segments of 1 second and TweetyNet’s output is the most likely label for each 2.7msec time bin in the recording.

#### Assuring the separation of syllable classes

To make sure that the syllable classes are well separated all the spectrograms of every instance of every syllable, as segmented in the previous section, were zero-padded to the same duration, pooled and divided into two equal sets. For each pair of syllable types, a support vector machine classifier was trained on half the data (the training set) and its error rate was calculated on the other half (the test set). These results are presented, for example, in Supplementary Figure 1 - 5.

#### Testing for within-class separation context-distinction by syllables acoustics

Apart from the clear between-class separation of different syllables for syllables that precede complex transitions we check the within-class distinction between contexts that affect the transition. To do that we use the parameters defined in Wohlgemuth et al., 2010^50^ and treat each syllable rendition as a point in an 8 dimensional space of normalized acoustic features. For a pair of syllable groups (different syllables or the same syllable in different contexts) we calculate the discriminability coefficient:

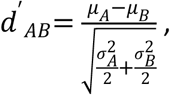

Where, *μ* _*A*_ −*μ* _*B*_ is the *L*_2_distance between the centers of the distributions and 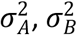 are the within-group distance variance from the centers. Supplementary Figure 1 - 7 demonstrates that all within-class d’ values are smaller than all between-class d’ values.

#### Identifying complex transitions

Complex transitions were identified by the length of the Markov chain, required to describe the outcome probabilities. These dependencies were found using a previously-described algorithm that extract the probabilistic suffix tree (PST^1^) for each transition (https://github.com/jmarkow/pst). Briefly, the tree is a directed graph in which each phrase type is a root node representing the first order (Markov) transition probabilities to downstream phrases, including the end of song. The pie chart in Supplementary Figure 1 - 4(i) shows such probabilities. Upstream nodes represent higher order Markov chains, 2^nd^ and 3^rd^ in Supplementary Figure 1 - 4(ii) and (iii) respectively, that are added sequentially if they significantly add information about the transition.

#### ROI selection, DF/F signal extraction and de-noising

Song-containing movies were converted to images by calculating, for each pixel, the maximal value across all frames. These ‘maximum projection images’ were then similarly used to create a daily maximum projection image and also concatenated to create a video. The daily maximum projection and song-wise maximum projection videos were used to select regions of interest (ROIs), purported single neurons, in which fluorescence fluctuated across songs.

ROIs were never smaller than the expected neuron size, did not overlap, and were restricted to connected shapes, rarely deviating from simple ellipses. Importantly, this selection method did not differentiate between sources of fixed and fluctuating fluorescence. The footprint of each ROI in the video frames was used to extract the time series, *F*(*t*) = *Σ*_(*x,y*) ∈*ROI*_ *V*(*x, y, t*), summing signal from all pixels within that ROI. Then, signals were converted to relative fluorescence changes, 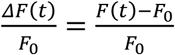 by defining *F* to be the 0.05 quantile.

The de-noised fluorescence, ((*Δf*/*f*_0_)_*denoised*_, is estimated from the relative fluorescence change using previously published modeling of the calcium concentration dynamics and the added noise process caused by the fluorescence measurement^46^

#### Seeking ROIs with sequence correlations

Since each ROI was sparsely active in very few phrase types we first sought ROIs that are active during a phrase type and then tested if it shows correlations to preceding or following phrase identities. We used the following scheme:

- The entire set of songs of each bird was used to calculate the 1^st^ order phrase transition probabilities, *P*_*ab*_ = *P*(*a* → *b*), for all phrases ‘a’, ‘b’.
- Phrase-type-active ROI was defined by requiring signal magnitude, 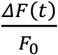 as defined in the previous section, to be larger than neighboring phrases and distinct from noise fluctuations (for each ROI and repeats of each phrase type, P):
  - The 0.9 quantile, *ΔFF*_90_, is taken as a measure of peak values – reducing outliers.
  - Signal and noise fluctuations are separated by fitting a 2-state hidden Markov model with Gaussian emission functions. The phrase-type-occupancy, *HMM*_*P*_, is the fraction of phrase ‘P’ repetitions that contained the signal state.
  - ROI past and future phrase type relationships were investigated if *ΔFF*_90_ > 0.1 (i.e. fluorescence fluctuation is large), *HMM*_*P*_ > 0.1 (i.e. the phrase type reliably carries signal), and *ΔFF*_90_ is larger than the same measure in the neighboring phrases (i.e. the phrase type carries a peak of signal value compared to its neighbors).
- Sequence-signal correlations were not investigated if fewer than N = 10 repeats contributed to the test.
- 1^st^ order relationships between the signal integral (summed across time bins in the phrase) and the upstream or downstream phrase identities were tested with 1-way ANOVA.
- 2^nd^ order relationships were tested between the signal integral and the identity of the 2^nd^ upstream (downstream) phrase identity for all intermediate phrase types that preceded (followed) the phrase-in-focus in ≥ 10% of the repeats (as indicated by the phrase transition matrix)
- Relations were discarded if the label, leading to the significant ANOVA, contained only one song.
- Data used for ANOVA tests is represented in supplementary figures by box plots marking the median (center line); upper and lower quartiles (box limits); extreme values (whiskers), and outliers (+ markers).
- The data were not tested for normality prior to performing ANOVA tests for individual neurons with the following reasoning:
  - Statistics textbooks suggest that violating the normality requirement is not expected to have a significant effect. For example, Howell, Statistical Methods in Psychology, Chapman & Hall, 4th Ed writes: “As we have seen, the analysis of variance is based on the assumptions of normality and homogeneity of variance. In practice, however, the analysis of variance is a robust statistical procedure, and the assumptions frequently can be violated with relatively minor effects. This is especially true for the normality assumption. For studies dealing with this problem, see Box (1953, 1954a, 1954b), Boneau (1960), Bradley (1964), and Grissom (2000).”
  - Carrying tests for normality will create a bias in our analyses. Each neuron tested for phrase sequence correlation is recorded in a different number of songs. Testing for normality will bias towards larger numbers of songs and against high-order correlations.
  - Nevertheless, we repeated the analyses in this manuscript with non-parametric one-way analysis of variance (Kruskal - Wallis). While fewer neurons pass the more stringent tests (~15% less), all the results in the manuscript remain the same. We include a summary of the non-parametric statistics as extended data.

#### Note

In this procedure, sparsely active ROIs or ROIs active in rare phrase types were not tested for sequence correlation. In the main body we reported that 18.2% of the entire set of ROIs showed sequence correlation. This percentage includes also ROIs that were not tested for sequence correlations. Out of the ROIs that were tested, about 30% had significant sequence correlations (23% and 10% showed 1^st^ and 2^nd^ order correlations)

#### Phrase specificity

The fractions of phrase repetitions, during which a ROI is ‘active’, *HMM*_*P*_, were also used to calculate the ROIs’ phrase specificity (in Figure 2):

- For each ROI, the fraction of activity in repetitions of each phrase was calculated separately.
- These measures were normalized and sorted in descending order.
- The number of phrase types accounting for 90% of the ROI’s activity was calculated.

#### Controlling for phrase durations and time-in-song confounds

In songs that contain a fixed phrase sequence, as in Figure 2d, we calculated the significance of the relation between *s* = Σ _*t*∈*p*_ (*Δf*/*f*_0_)_*denoised*_, an integral of the signal during one phrase in the sequence, the target phrase ‘p’, and the identity of an upstream phrase that changes from song to song using a 1-way ANOVA. This relation can be carried by several confounding variables:

- The duration of the target phrase.
- The relative timing of intermediate phrase edges, between the changing phrase and the target phrase.
- The absolute time-in-song of the target phrase.

In Supplementary Figure 2 - 7c we account for these variables by first calculating the residuals of a multivariate linear regression (a general linear model, or GLM) between those variables and *s t*, and then using 1-way ANOVA to test the relation of the residuals and the upstream or downstream phrase identity.

#### Contrasting influence of preceding and following phrases on neural activity (Supplementary Figure 3 - 3)

For neurons with significant sequence correlations (1-way ANOVA described above) we adopted a method agnostic to correlation order (1^st^ or higher, as defined above) and direction (past or future). We used multi-way ANOVA to test the effect of the identity of the immediately preceding and immediately following phrase types on the neural signal (*s* = Σ∈*p Δf*/*f*_0_)*denoised*). Using a threshold at p = 0.05 we compare the fractions of sequence-correlated ROIs influenced by past phrases, future phrases, or both. This comparison is also carried separately for ROIs active in complex transitions or outside of complex transitions (panels b,c).

#### Testing if sequence correlated neurons prefer one or more song contexts (Figure 4c)

For neurons with significant sequence correlations (1-way ANOVA described above) we used Tukey’s post-hoc analysis to determine if this sequence correlation results from a significant single preferred context or significant several preferred contexts. A neuron was declared ‘single context preferring’ if the mean signal in only that context was larger than all others (Tukey’s p<0.001). A neuron was declared as having preference to more than a single past context if the mean signal following several contexts was larger than another context (Tukey’s p<0.001). Since the post-hoc test uses a subset of the songs it is weaker than the 1-way ANOVA and some neurons do not show a clear preference to one context or more but still have sequence correlation (gray in Figure 4c)

#### Maximum projection images for comparing context-dependent signals

##### Maximum fluorescence images

In songs that contain a fixed phrase sequence and a variable context element, such as a preceding phrase identity, maximum projection images are created, as above, but using only video frames from the target phrase (e.g. the pink phrase in Figure 2d). Then, the sets of maximum projection images in each context (e.g. identity of upstream phrase) are averaged, assigned orthogonal color maps (e.g. red and cyan in Supplementary Figure 2 - 3) and overlaid. Consequentially, regions of the imaging plane that have no sequence preference will be closer to gray scale, while ROIs with sequence preference will be colored. In Supplementary Figure 2 - 3 and Supplementary Figure 4 1 we used a sigmoidal transform of the color saturation to amplify the contrast between color and gray scale without changing the sequence preference information. Additionally, to show that pixels in the ROI are biased towards the same context preference, the above context-averaged maximum projection images are subtracted and pseudo-colored (insets in Supplementary Figure 2 - 3)

##### De-noised Maximum projection images (Figure 3a)

The above maximum projection images show the fluorescence signal, including background levels that are typical to single-photon microscopy. To emphasize context-dependent ROIs we de-noised the fluorescence videos using the previously-published algorithm, CNMFE^46^, and created maximum projection images, as above, from the background-subtracted videos. The preceding context preferring ROIs from this estimation algorithm (Figure 3a) completely overlap with the manually defined ROIs, used to extract signal rasters (Figure 3b). Supplementary Figure 3 - 3 replicates Figure 3a without the de-noising algorithm and shows that the same ROIs report the same context dependence. Supplementary video 8 shows all the de-noised video data, used to create Figure 3a.

#### Label prediction from clustered network states

The signal integral during a target phrase, pink in Figure 4d, is used to create network states – vectors, composed of signals from 4 jointly-recorded ROIs. The averages of the vectors, belonging to the contexts defined by the 1^st^ upstream (or downstream) phrase label define label-centroids. Then, labels of individual songs are assigned to the nearest neighboring centroid (Euclidean).

#### Bootstrapping mutual information in limited song numbers

To overcome the errors introduced by limited number of songs in Figure 4d, the mutual information between the network state and the identity of the 1^st^ upstream (or downstream) phrase is estimated in a bootstrapping permutation process as follows:

- Sub-sampling 3 out of 4 ROIs in each permutation and converting their signal to binary values by thresholding the signal integral.
- Reducing the number of phrase labels by merging. Specifically, in Figure 4, the least common label in downstream states is randomly merged with one of the other labels. In the upstream labels, the least common label is merged after a random division of the other 4 labels, to form 2 groups of 2.
- The mutual information measures are then calculated for each one of the 48 possible state spaces – leading to the scatter Figure 4e. The 0.95 quantile level of the null hypothesis is created by randomly shuffling each variable to create a 1000 surrogate data sets and repeating the measures.
- The shuffled set is used to create a sample distribution and calculate the significance of the differences in Figure 4e using a z-test with the sample mean and standard deviation.

