## Supplemental analysis 1 for "Hidden neural states underlie canary song syntax"

**Extended data – non-parametric statistical analysis for results in main manuscript**

Several analyses in the main manuscript “Hidden neural states underlie canary song syntax” are based on 1-way ANOVA tests. While statistics textbooks suggest that the requirement for normally distributed data can be relaxed without significant effects (c.f. quotes and citations in the Methods section) we repeated the manuscript’s analyses with the non-parametric 1-way ANOVA (Kruskal Wallis). While fewer neurons pass the more stringent tests (~15% less), these analyses recapitulated all the findings in the manuscript. This section summarizes the statistics and results of the non-parametric tests.

As in the main text, in the following, the number of categories is small and we report the 2^nd^ number of degrees of freedom.

**Statistics in Supplementary figure 2-2 showing the sequence correlations of the example neuron in Figure 2d**:

Naïve: $\chi^{2}\left( 35 \right)=27.37, p<5{\cdot10}^{-5}$

Removing dependency on phrase duration: $\chi^{2}\left( 35 \right)=13.64, p<0.02$

Removing dependency all phrase edges in the sequence: $\chi^{2}\left( 35 \right)=13.03, p<0.025$

Removing dependency on all edges and the global time in song: $\chi^{2}\left( 35 \right)=12.75, p<0.03$

**Statistics in Supplementary figure 2-3 panels**:

a. $\chi^{2}\left( 27 \right)=16.31, p<0.0011$
b. $\chi^{2}\left( 14 \right)=4.26, p<0.04$
c. $\chi^{2}\left( 19 \right)=11.11, p<0.012$
d. $\chi^{2}\left( 21 \right)=5.16, p<0.024$
e. $\chi^{2}\left( 14 \right)=6.73, p<0.01$
f. $\chi^{2}\left( 28 \right)=10.08, p<0.0016$

**Fractions of daily annotated ROIs with significant sequence correlations**:

Total: 18.22%. 1^st^ order: 15.72%, $\geq$2^nd^ order: 4.2%.

**Statistics in Supplementary figure 2-7d**:

Accounting for all confounding variables: 51.3% of 1^st^ order tests remain significant and 29.7% of $\geq$2^nd^ order tests remain significant.

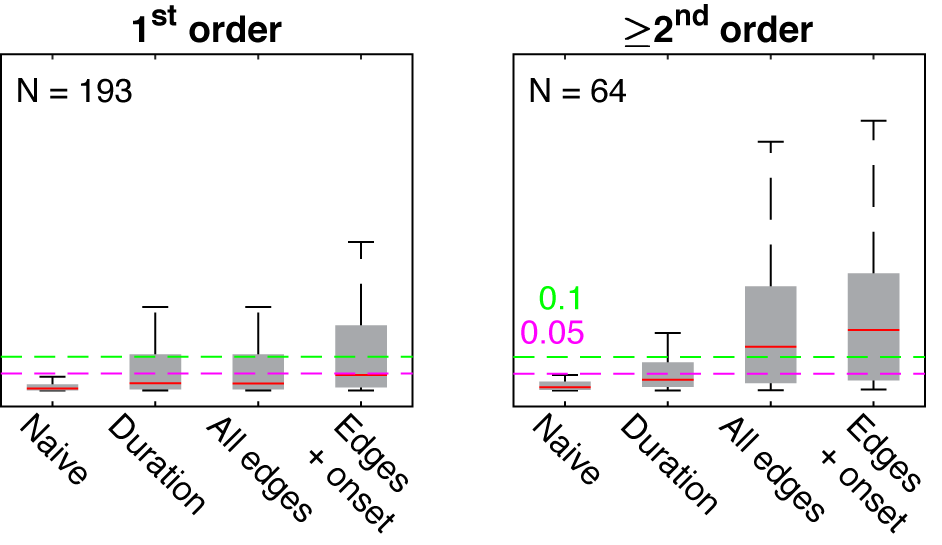

**Statistic in Figure 2e showing more neurons encoding past events**:

With non-parametric test the bias remains and slightly increases for high order sequence correlations. 67.9% encode the past in 1^st^ order relations and 65.6% in $\geq$2^nd^ order relations (all highly significant biases)

**Statistical tests in Figure 3**:

ROI #21 during first phrase: $\chi^{2}\left( 13 \right)=10.12, p<0.0016$

ROI #45 during first phrase: $\chi^{2}\left( 13 \right)=4.5, p<0.034$

ROI #50 during second phrase: $\chi^{2}\left( 13 \right)=10.12, p<0.0016$

ROI #36 during third phrase: $\chi^{2}\left( 13 \right)=5.56, p<0.019$

ROI #21 during last phrase: $\chi^{2}\left( 13 \right)=6.72, p<0.0016$

ROI #45 during last phrase: $\chi^{2}\left( 13 \right)=8.68, p<0.0035$

In total 68.3% of sequence-correlated neurons are found in complex transitions.

**Statistical tests in Supplementary Figure 4-1 panels** **supporting Figure 4**:

a. $\chi^{2}\left( 36 \right)=25.44, p<4.1\cdot{10}^{-5}$
b. $\chi^{2}\left( 13 \right)=14.24, p<0.015$
c. $\chi^{2}\left( 40 \right)=28.66, p<2.65\cdot{10}^{-6}$
d. $\chi^{2}\left( 41 \right)=20.37, p<0.00011$
e. $\chi^{2}\left( 23 \right)=20.23, p<0.0026$
