## Supplemental analysis 2 for "Hidden neural states underlie canary song syntax"

### Appendix A - Supplementary analysis of overlapping ROIs

In single photon calcium imaging, especially with head-mounted microscopes, the treatment of regions of interest (ROIs) that overlap between days is controversial (Liberti et al 2016, Katlowitz et al 2018). In this study, canary neural signals were acquired with chronically implanted mini-microscopes and it is plausible to assume that the same sources appeared in multiple days. Due to the controversy in the literature we chose to treat ROIs independently each day. This choice is justified by two reasons. First, the numbers of identified ROIs per day often exceeded 30 (peak values exceeding 70) and these sample sizes should robustly approach the mean of the per-day distributions for the tests we carried. Second, grouping ROIs across days can possibly introduce uncontrolled biases.

Never the less, we also carried an additional analysis with the following underlying assumptions:

- ROIs that overlapped in at least 25% (of the smaller ROI) across consecutive sessions were suspected as the same source.
- Suspected ROIs that showed the same significant sequence preference in at least one context (as in the methods section of the main manuscript) were considered as candidates for a grouped representation. This is a permissive choice since ROIs could be active in more than a single context. In doing so we only reduce the numbers of ROIs with sequence correlations and the statistical power of subsequent tests.
- In the following tests for the population of sequence correlated ROIs we treated suspected ROIs as a single grouped representation in each test.

**There are more sequence correlates of past phrases than future phrases**

We repeated the analysis in figure 2e of the main manuscript (Figure 1b, below). With grouping ROIs across days the numbers of representations decrease by about a half. With this massive grouping, the main observation remains the same. There are more representations of past phrases of both first and second (or higher) orders (Figure 1a, below). The difference remains highly significant in first order representation but becomes statistically insignificant in higher order representations. This difference does not change the discussion in the main manuscript.

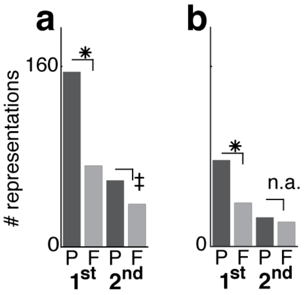


Figure 1 | **Grouping ROIs does not account for the bias in representing more past context than future context. a.** Figure 2e of the main manuscript shows more individual ROI correlates to past events than to future events. Numbers of significant correlations with adjacent (1st order, 2 left bars) and non-adjacent (≥2nd order, 2 right bars) phrases. The correlations are separated by phrases that precede (P) or follow (F) the phrase, during which the signal is integrated. Binomial z-test evaluate significant differences (✳: $p < 1e-10$, ‡: $p < 0.002$). **b.** The same ROIs after grouping have more representation of past events. This difference is significant for the 1$^{st}$ order representations. (✳: $p < 1e-6$, n.a.: $p > 0.1$, binomial z-tests).

**Sequence correlates are predominately found in complex transitions**

As in the previous section, grouping ROIs across days does not change the finding in the main manuscript (Figure 3c of the main manuscript, Figure 2a below). Most of the grouped correlates are found during complex transitions (Figure 2b below).

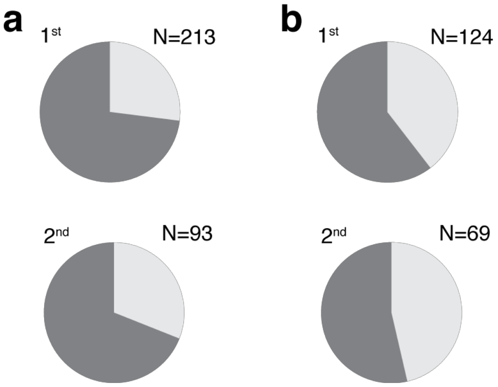


Figure 2 | **Grouped sequence representations are also predominately found during complex transitions. a.** Figure 3c of the main manuscript shows the fraction of sequence-correlated ROIs found in complex transitions. Pie charts separate 1st order and higher order (≥2nd) sequence correlations. Dark grey summarizes the total fraction for two birds. **b.** The same ROIs after grouping are also mostly found in complex transitions.
